# Comprehensive analysis of disease pathology in immunocompetent and immunocompromised hamster models of SARS-CoV-2 infection

**DOI:** 10.1101/2022.01.07.475406

**Authors:** Santhamani Ramasamy, Afsal Kolloli, Ranjeet Kumar, Seema Husain, Patricia Soteropoulos, Theresa L. Chang, Selvakumar Subbian

## Abstract

The pathogenesis of SARS-CoV-2 in the context of a specific immunological niche is not fully understood. Here, we used a golden Syrian hamster model to systematically evaluate the kinetics of host response to SARS-CoV-2 infection, following disease pathology, viral loads, antibody responses, and inflammatory cytokine expression in multiple organs. The kinetics of SARS-CoV-2 pathogenesis and genomewide lung transcriptome was also compared between immunocompetent and immunocompromised hamsters. We observed that the body weight loss was proportional to the SARS-CoV-2 infectious dose and lasted for a short time only in immunocompetent hamsters. Body weight loss was more prominent and prolonged in infected immunocompromised hamsters. While the kinetics of viral replication and peak live viral loads were not significantly different at low and high infectious doses (LD and HD), the HD-infected immunocompetent animals developed severe lung disease pathology. The immunocompetent animals cleared the live virus in all tested tissues by 12 days post-infection and generated a robust serum antibody response. In contrast, immunocompromised hamsters mounted an inadequate SARS-CoV-2 neutralizing antibody response, and the virus was detected in the pulmonary and multiple extrapulmonary organs until 16 days post-infection. These hamsters also had prolonged moderate inflammation with severe bronchiolar-alveolar hyperplasia/metaplasia. Consistent with the difference in disease presentation, distinct changes in the expression of inflammation and immune cell response pathways and network genes were seen in the lungs of infected immunocompetent and immunocompromised animals. This study highlights the interplay between the kinetics of viral replication and the dynamics of SARS-CoV-2 pathogenesis at organ-level niches and maps how COVID-19 symptoms vary in different immune contexts. Together, our data suggest that the histopathological manifestations caused by progressive SARS-CoV-2 infection may be a better predictor of COVID-19 severity than individual measures of viral load, antibody response, and cytokine storm at the systemic or local (lungs) levels in the immunocompetent and immunocompromised hosts.

## 1. INTRODUCTION

Coronavirus disease-2019 (COVID-19), caused by the severe acute respiratory syndrome coronavirus-2 (SARS-CoV-2), continues to be a significant global health concern. To date, at least 250 million cases and 5 million deaths have been confirmed worldwide^1^. COVID-19 pathology is associated with host immune alterations, which begin during the early stages of infection and lead to uncontrolled SARS-CoV-2 replication. The resulting hyper-inflammation progresses to acute respiratory distress syndrome (ARDS) and ultimately death in the most severe cases^2, 3^.COVID-19 patients with delayed virus clearance (>20 days) were at higher risk of progression to ARDS and death^4, 5^. Further, preexisting host immunosuppression is a significant risk factor for delayed virus clearance and compromised antibody response to SARS-CoV-2. For example, patients immunocompromised by multiple myeloma died of ARDS following SARS-CoV-2 infection due to their inability to produce antibodies and clear the virus^6^. However, treatment with cyclosporine, an immunosuppressive agent, did not affect viral clearance in a kidney transplant patient infected with SARS-CoV-2 ^6^. Therefore, the intricate association between immunosuppression and its impact on the progression of SARS-CoV-2 infection remains unclear.

Though the degree of pulmonary pathology, including pneumonia and ARDS, are strong predictors of the severity of COVID-19, the disease is a multisystem disorder that can compromise neuronal, cardiovascular, gastrointestinal, urogenital, endocrine, and other systems ^7^. Therefore, it is imperative to analyze the extent of disease pathology in the extrapulmonary system to understand pathological complications, which are currently poorly defined^8^. A key challenge to identifying and characterizing pathology caused by SARS-CoV-2 in various tissues is biosafety-based restrictions on autopsies of patients who died of COVID-19 ^9^. The reports on COVID-19 pathology in animal models are currently confined mainly to the respiratory tract, and the systematic, longitudinal analysis of extrapulmonary pathology is scant^10–12^.

Antibody-mediated immune response, elicited in the host upon vaccination and exposure to SARS-CoV-2, can protect against the progression of infection. Further, the extent of protection correlates with the viral-neutralizing antibody titer in patient sera ^13^. It has been shown that immunocompromised COVID-19 patients lack a strong antibody response against SARS-CoV-2, which would explain severe disease in these individuals^6^. However, the effect of reducing hyper-inflammation by steroids and other immunosuppressive treatments on SARS-CoV-2 infection is not fully understood. Since immunocompromised hosts are more vulnerable to severe COVID-19, it is crucial to understand the impact of cyclophosphamide treatment-mediated immunosuppression on the progression of SARS-CoV-2 infection.

Previous studies show that SARS-CoV-2 infected Golden Syrian hamsters are reliable and predictable animal models that can recapitulate the pulmonary pathological manifestations of COVID-19 in humans^14–17^. These reports show that following intranasal SARS-CoV-2 inoculation, high viral load and disease pathology were mainly localized to the trachea and lungs, with interstitial pneumonia being the main hallmark of severe disease ^14–17^. At the cellular level, a relatively higher proportion of B cells and neutrophils were noted in the lungs of COVID-19 patients. In contrast, fewer macrophages with concomitantly higher NK cell numbers were observed in patients recovered from SARS-CoV-2 infection ^18^. Further, immunoglobulin production, humoral immune response, cytokine signaling, interferon signaling, T cell activation signaling pathway genes were upregulated during progressive SARS-CoV-2 infection in immunocompetent individuals^18, 19^. Consistent with these observations in humans, the inflammatory pathway, type I IFN, and IFNG were upregulated in the lungs of hamsters early during SARS-CoV-2 infection ^20^. However, systematic pathologic evaluation of various internal organs impacted by SARS-CoV-2, and the association between disease pathology and viral replication kinetics, were not thoroughly analyzed. Importantly, the immunopathology of SARS-CoV-2 infection in an immunocompromised hamster model that mimics COVID-19 in individuals with health conditions such as cancer and autoimmune disease has not been examined previously.

This study reports the systematic analysis of disease severity across organs in a hamster model of pulmonary SARS-CoV-2 infection. We analyzed the kinetics of viral replication, antibody response, cytokine storm marker expression, and disease pathology in pulmonary and extrapulmonary tissues of immunocompetent and immunocompromised hamsters infected intranasally with a low or high dose of a virulent strain of SARS-CoV-2. Further, using genomewide RNAseq analysis of the lungs, we determined the differential immune correlates of disease pathogenesis between immunocompetent and immunocompromised hamsters during SARS-CoV-2 infection. Our data suggest that the disease pathology in the pulmonary and extrapulmonary systems are the key and prominent indicators of the severity of progressive SARS-CoV-2 infection than the serum antibody response and viral load in internal organs. The differential expression of network and pathway genes between immunocompetent and immunocompromised hamsters corroborate the underpinning differential SARS-CoV-2 pathogenesis in these animals.

## 2. RESULTS

### 2.1. Distinct disease progression in SARS-CoV-2 infected immunocompetent and immunocompromised hamsters

To understand disease pathology upon pulmonary SARS-CoV-2 infection, hamsters were intranasally inoculated with a low dose (LD; 10^2.5^ PFU) or high-dose (HD; 10^6^ PFU) inoculum of virus (see methods). Following intranasal infection, both the LD and HD-infected hamsters showed a reduction in body weight from the day of infection until six days post-infection (dpi). The animals then gradually gained weight from 8 to 16 dpi, suggesting that they experienced inappetence or anorexia during the early stages of the disease, as determined by food/water intake. A maximum mean weight loss of 2.63 and 8.4%, respectively, was observed in LD and HD infection groups at 6 dpi (Figure 1a). To model immunosuppression, a group of hamsters was treated with cyclophosphamide before and during LD intranasal SARS-CoV-2 infection (see Methods). In contrast to the immunocompetent animals, the immunocompromised (CP-LD) hamsters showed slow weight loss up to 8 dpi from the day of infection. These animals also showed a slow-weight gain between 8 and 16 dpi (Figure 1a). The maximum mean weight loss observed was 7.67 % at 8 dpi. However, the mean body weight on day 16 was only 2.8 % higher than the mean weight at infection (Figure 1a).

**Figure 1:**
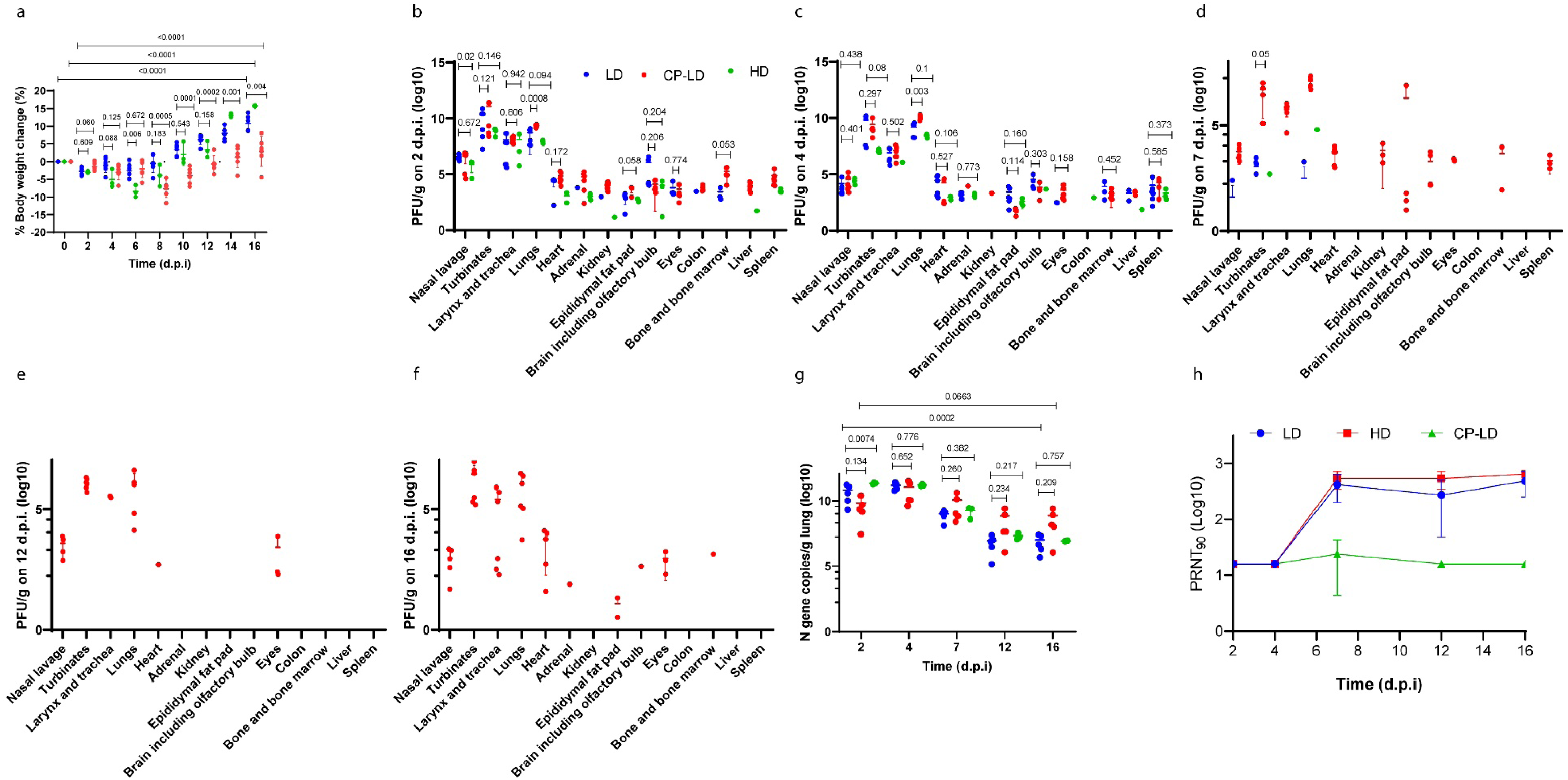
COVID-19 disease in SARS-CoV-2 infected hamsters. Comparison of body weight change **(a)** in LD (10^2.5^ PFU, n=30), HD (10^5^ PFU, n=15) and CP-LD (immunocompromised; 10^2.5^ PFU, n=30) SARS-CoV-2 infected hamsters over 2-16 dpi. Median weight change (g) over 2-16 dpi, compared to weight at the time of infection. Comparison of viral load **(b-f)** in SARS-CoV-2 infected hamsters (LD vs. HD, and LD vs. CP-LD) on 2-16 dpi, expressed as PFU/g of tissues. The expression of SARS-CoV-2 N gene copies/g of lungs over 2-16 dpi in LD, HD, and CP-LD infected hamsters **(g)**. Kinetics of plaque reduction neutralization titer in hamster sera collected at 2-16 dpi **(h)**. statistical analysis was performed by two-sided Welch’s t-test **(a-d, g)** and one-way ANOVA **(a, g)**. p values are indicated above the plots. Statistical analysis was performed by comparing the three or more data points at any organs **(b-f)**. Data represent mean ±s.d. (**a-h).** The blue, green, and red indicate LD, HD, and CP-LD infected hamsters, respectively, and each dot represents data from an individual hamster. LD and CP-LD; n=6, HD, n=3 per time point.

### 2.2. Viral burden in immunocompetent hamsters infected with a low or high dose inoculum

We performed a plaque-forming unit (PFU) assay on tissue homogenates to determine replicating viral load in various tissues. In both LD and HD-infected hamsters, the highest viral burden was detected in the lungs, turbinate, and larynx/trachea at 2 dpi (Figure 1b). The viral load in these organs did not significantly change between 2 and 4 dpi. At this timepoint, infectious viral particles were also detected in the heart, adrenal gland, epididymal fat pad, brain (including olfactory bulb), and eyes of both LD and HD infected animals (Figure 1b, c). However, at both 2 and 4 dpi, the viral load in these extrapulmonary organs was about 3-4 logs, significantly lower than in the lungs (Figure 1b, c).

In the LD-infected hamsters, viruses were detected in the colon, kidney, and adrenal glands of 1 out of 6 animals at 2 and 4 dpi in the liver and spleen. In contrast, at 4 dpi, none of the hamsters had viruses in the colon and kidney, and 3 had viruses in the adrenal glands (Figure 1b, c, Table 1). Four out of 6 hamsters at 2 dpi and all 6dpi at 4 dpi had infectious viral particles in the heart. However, the viral load in the extrapulmonary organs was not significantly different between 2 and 4 dpi (Figure 1b and 1c). Four out of 6 hamsters at 7 dpi showed viral load in the nasal turbinate, while one animal each showed viruses in the lungs and nasal lavage (Figure 1d, Table 1). The viral load in these tissues was significantly lower at 7 dpi than at 2 and 4 dpi (Figure 1b-d).

**Table 1:**
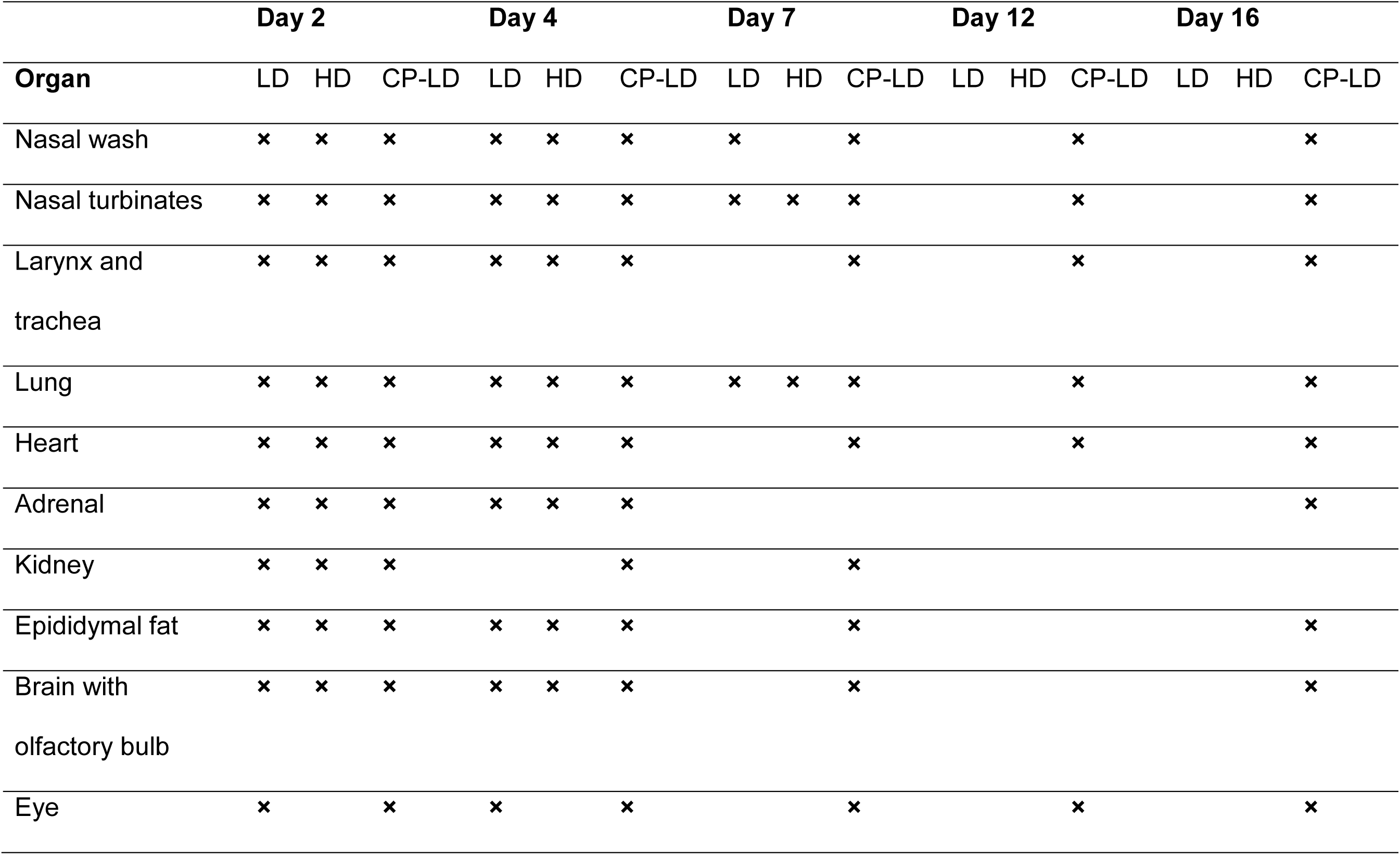
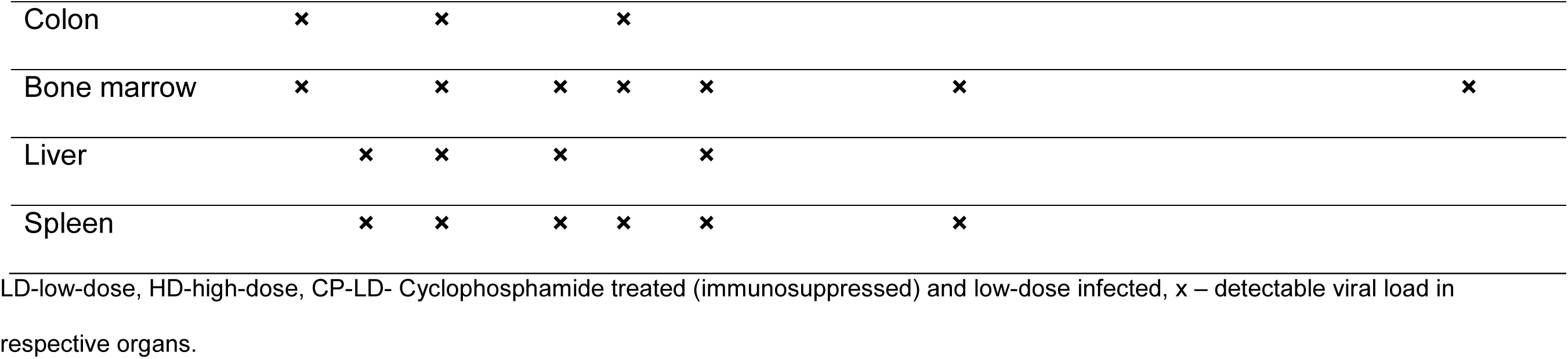
The detectable SARS-CoV-2 in infected hamster organs at different time points.

Following HD infection, peak viral load was detected at 2 and 4 dpi. Two out of 3 hamsters had viruses in the heart, spleen, adrenal, epididymal fat, and brain at 4 dpi. At this time, 1 out of 3 hamsters showed viral load in the liver, kidney, and colon (Figure 1c). At 7 dpi, the infectious virions were detectable in the lungs and turbinates of 1 out of 3 hamsters (Figure 1d). The viral loads in the lung, nasal lavage, nasal turbinates, larynx/trachea, heart, epididymal fat, brain were not significantly different between LD and HD at 2 and 4 dpi (Figure 1b, c). To summarize, the viral burden in pulmonary and extrapulmonary organs was not significantly different between LD and HD SARS-CoV-2 infected immunocompetent hamsters on 2, 4, 7, 12, and 16 dpi.

### 2.3. Disparate tissue viral burden between immunocompetent and immunocompromised hamsters infected with SARS-CoV-2

Next, we examined the effect of immune suppression on SARS-CoV-2 replication and time to viral clearance in tissues. Hamsters were treated with cyclophosphamide, infected with LD inoculum (CP-LD), and infectious virus was determined in the tissue homogenates. In contrast to LD-infected immunocompetent hamsters, the immunocompromised animals had infectious SARS-CoV-2 virions in their respiratory tract from 2 dpi until 16 dpi (Figure 1b-f). In these animals, the viral load in nasal lavage gradually decreased from 2 dpi until 7 dpi and stabilized at similar levels up to 16dpi (Figure 1b-f). Viral load peaked in the lungs at 4 dpi in the immunocompromised hamsters (CP-LD) (Figure 1c), then gradually declined between 7 dpi to 12 dpi (Figure 1d, e). A similar viral load was observed between 12 and 16 dpi (Figure 1e and 1f). In contrast, peak viral load was noted at 2 dpi in the nasal turbinates and larynx/trachea, which gradually declined at 7dpi (Figure 1b, d). The viral load was not significantly different in these organs at 7, 12, and 16 dpi. The infectious virus in the heart was detectable at 2, 4, and 7dpi that declined at 12dpi (1 out of 6 animals had the virus), before rebounding at 16 dpi (5 out of 6 had a detectable virus) (Figure 1b-f). All the immunocompromised and infected hamsters had infectious viruses in the adrenal gland, brain, eye, colon, bone marrow, liver and spleen; 5 out of 6 animals in kidneys and 4 out of 6 in epididymal fat at 2 dpi (Figure 1b). The mean viral load in the adrenal gland, bone marrow, liver, and colon declined gradually from 2 until no live virus was detected in 7dpi (Figure 1b-d). At 12 dpi, the virus was detected only in the heart (1 out of 6), eyes (3/6), and respiratory system (Figure 1e). Interestingly, infectious viruses were detected in the heart, adrenal gland, epididymal fat, brain, eye, and bone marrow of some immunocompromised and infected hamsters at 16 dpi (Figure 1f).

No significant difference in the viral load was observed in the nasal lavage, turbinates, larynx/trachea, heart, adrenal gland, brain, eyes, and bone marrow between immunocompetent (LD) and immunosuppressed (CP-LD) animals, at 2 and 4 dpi. However, a significantly higher viral load was noted in the lungs of infected immunocompromised hamsters at 2 and 4 dpi and in nasal lavage of immunocompetent hamsters at 2 dpi. Similarly, the live virus was detected in the liver and spleen of only immunocompromised hamsters at 2 dpi and in the kidneys at 4 dpi. Replicating viral load was observed in the heart, adrenal, epididymal fat pad, brain, eyes, and bone marrow in these animals at 7 and 16 dpi (Figure 1b-f).

The total viral burden in the lung was also determined by measuring the SARS-CoV-2 N gene (+ and – strand RNA) transcripts, which were detected in the lungs of immunocompromised and immunocompetent infected hamsters (both LD and HD) from day 2 to 16 dpi. A peak in viral RNA load was observed in LD and HD groups at 2 and 4 dpi, although the viral RNA load was not significantly different between these two groups at 2, 4, 7, 12 and 16 dpi (Figure 1g). Interestingly, although more infectious viruses were detected in the infected immunocompromised hamster lungs from day 2 to 16 dpi, the viral RNA load was not significantly different from the immunocompetent (LD) group (Figure 1g).

### 2.4. Heterogeneity in neutralizing antibody titer in SARS-CoV-2 infected immunocompetent and immunocompromised hamsters

Sera from SARS-CoV-2 infected hamsters collected at 2, 4, 7, 12, and 16 dpi were analyzed for the virus-neutralizing antibodies. The plaque reduction neutralization test (PRNT) revealed that both LD and HD SARS-CoV-2 infected hamsters elicited detectable virus-neutralizing antibodies at 7 dpi. At 7 and 16 dpi, LD infected hamsters showed 90 % in PRNT neutralization (PRNT_90_) at 1:160 to 1:640 dilutions, respectively. The HD-infected hamsters showed PRNT_90_ at 1:360 to 1:640 serum dilutions at these time points (Figure 1h). Notably, the sera from immunocompromised hamsters showed PRNT_90_ at <1:16 dilution except one hamster serum at 7 dpi, which showed a PRNT_90_ at 1:64 dilutions (Figure 1h). These observations suggest that SARS-CoV-2 neutralizing antibody production was reduced in immunocompromised hamsters. Further, the presence of viable SARS-CoV-2 until 16 dpi in the infected immunocompromised animals, as well as a significant reduction or absence of infectious viruses starting at 7dpi in LD and HD infected immunocompetent hamsters indicate that the extent and rapidity of SARS-CoV-2 clearance are associated with the levels of production of virus-neutralizing antibodies at the systemic level.

### 2.5. Severity of pulmonary/bronchiolar pathology and thrombosis in SARS-CoV-2 infected immunocompetent and immunocompromised hamsters

Histopathological examination of SARS-CoV-2 infected lung samples at 4, 7 and 16 dpi were performed on hematoxylin and eosin-stained sections (Figure 2a-x). In both LD (Figure 2a-h) and HD (Figure 2i-p) groups, the lungs showed multifocal to diffuse infiltration of mononuclear cells in the interstitial spaces with the maximum infiltration on 4 (Figure 2a,b, e, f, i, j, m, n) and 7 dpi (Figure 2c, d, g, h, k, l, o, p). The inflammatory cellular infiltrations obliterated the alveoli and resulted in alveolar collapse (Figure 2b, c, j, k). A lower degree of inflammatory cell infiltration was noted in the immunocompromised animals at 4 and 7 dpi (Figure 2r, t) compared to immunocompetent hamsters infected with LD SARS-CoV-2 (Figure 2b, c, s, w).The pulmonary parenchyma of immunocompetent hamsters infected with LD SARS-CoV-2 showed moderate levels of mononuclear cells infiltration in the interstitium (Figure 2a, b, e and Table 2), congestion of capillaries in the alveolar wall (Figure 2a, b), mild bronchiolar epithelial hyperplasia (Figure 2b), bronchiolitis with lymphocytes infiltration and necrotic bronchiolar epithelial cells in the bronchiolar lumen (Figure 2b, f) at 4 dpi. In these animals, the presence of foamy macrophages and interstitial edema was prominent at 7 dpi. (Figure 2c, g).

**Figure 2:**
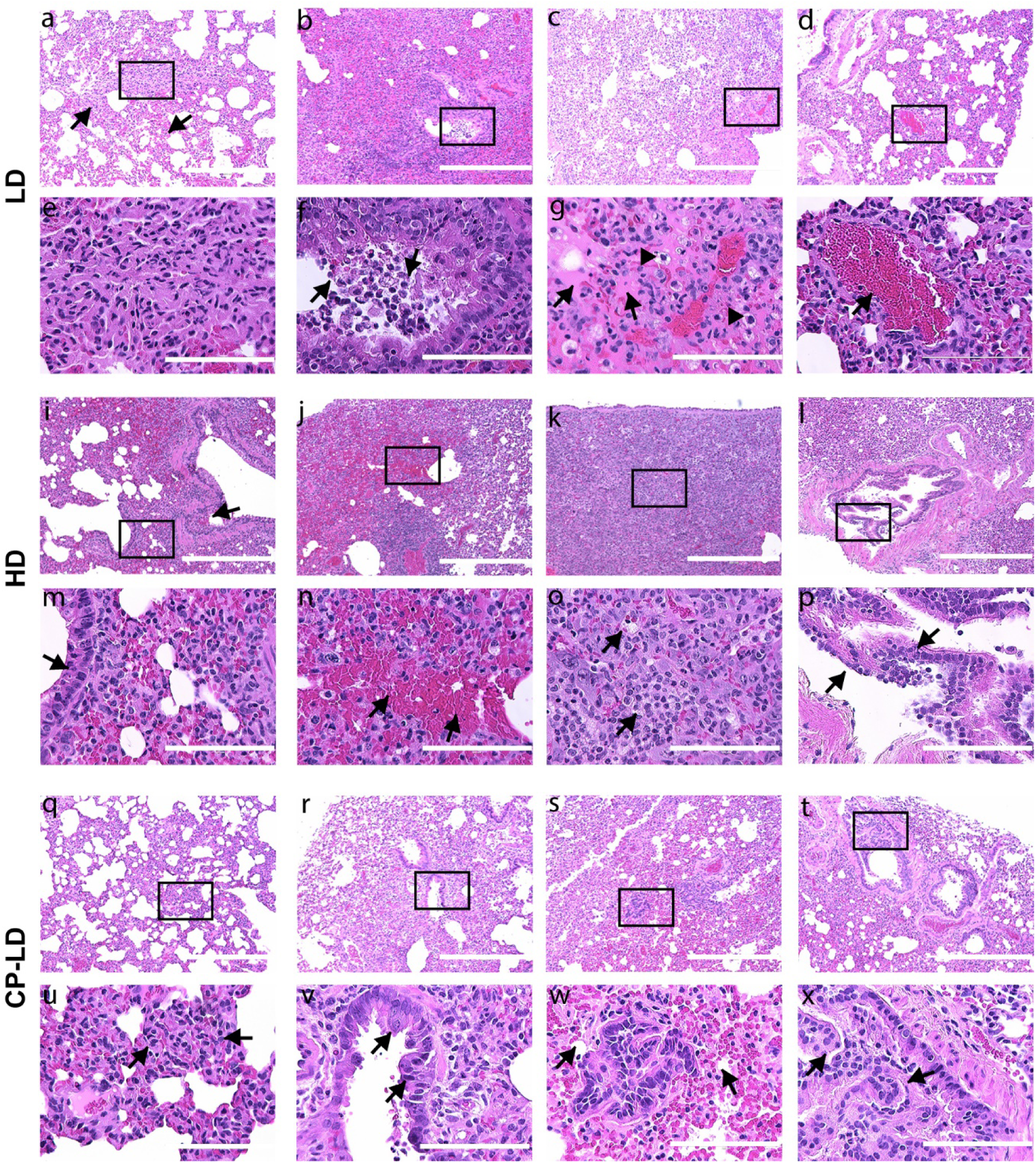
Interstitial pneumonia, bronchioalveolar and vascular pathology in SARS-CoV-2 infected hamsters. Histopathological analysis of LD, HD, CP-LD infected hamster lungs revealed infiltration of mononuclear cells in the interstitium **(a-h;** *arrows in* **a)**, in the bronchiolar lumen **(b**, **f;** *arrows in* **f)**, alveolar and interstitial edema **(c, g;** *arrows in* **g)**, foamy macrophages (*arrows heads in* **g)** and early thrombus **(c**, **g**, **d**, **h;** *arrow in* **h)**, interstitial/alveolar capillary congestion **(b)** of LD infected hamsters. Bronchiolar hyperplasia **(i**, **m**, **l**, **p;** *arrow in* **i)**, alveolar epithelial metaplasia **(I**, **m;** *arrow in* **m)**, interstitial hemorrhages **(j**, **n;** *arrows in* **n)**, and severe infiltration of inflammatory cells in the alveoli and interstitium caused the obliteration of alveoli in HD infected hamsters. Foamy macrophages **(j**, **n**, **k**, **o;** *arrows in* **o)**, bronchiolar epithelial hyperplasia and detachment, bronchiolitis, bronchiolar smooth muscle hyperplasia **(l**, **p;** *arrows in* **p)** were also noted in HD infected hamsters. Moderate infiltration of mononuclear cells and congestion of the interstitium (**q-x;** *arrows in* **u**), multiple alveolar epithelial metaplasia (*arrows in* **v)**, and obliteration of alveoli **(r**, **v**, **s**, **w)**, severe bronchiolar epithelial hyperplasia, and obliteration of lumen **(t**, **x)** were observed in the immunosuppressed infected (CP-LD) hamsters. Images **a-d**, **i-l**, **q-t** are 100x magnifications, and **e-f**, **m-p**, **u-x** are 400x magnifications (marked in 100x images). Scale bar represents 100 µ **(e-f**, **m-p**, **u-x)** or 400 µ **(a-d**, **i-l**, **q-t)**. Note: panels **a**, **b**, **e**, **f**, **i**, **j**, **m**, **n**, **q**, **u**, **t**, **x** are 4 dpi, **c**, **g**, **k**, **o**, **r**, **v** are 7 dpi, and **d**, **h**, **j**, **p**, **s**, **w** are 16 dpi.

**Table 2:**
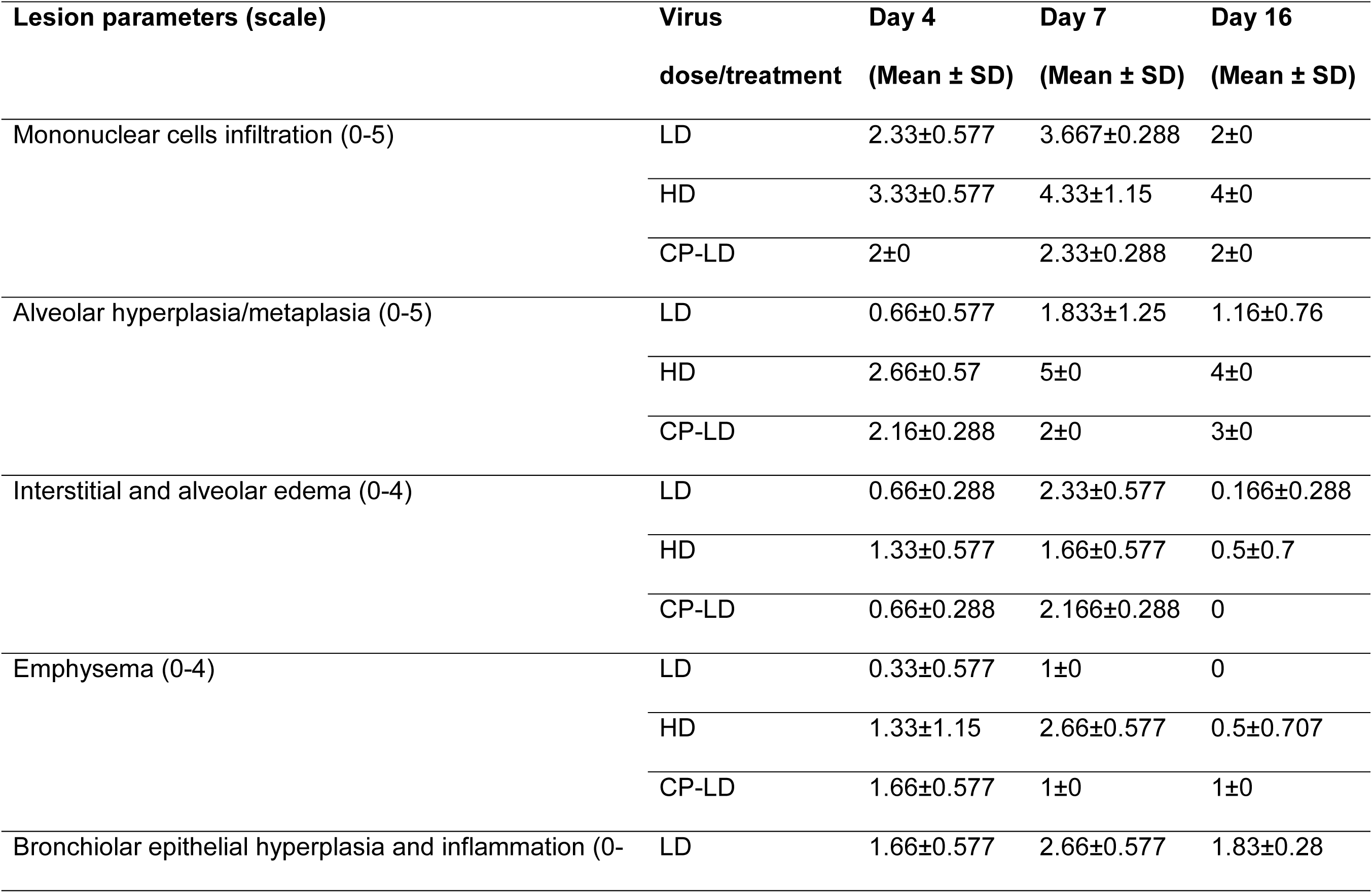
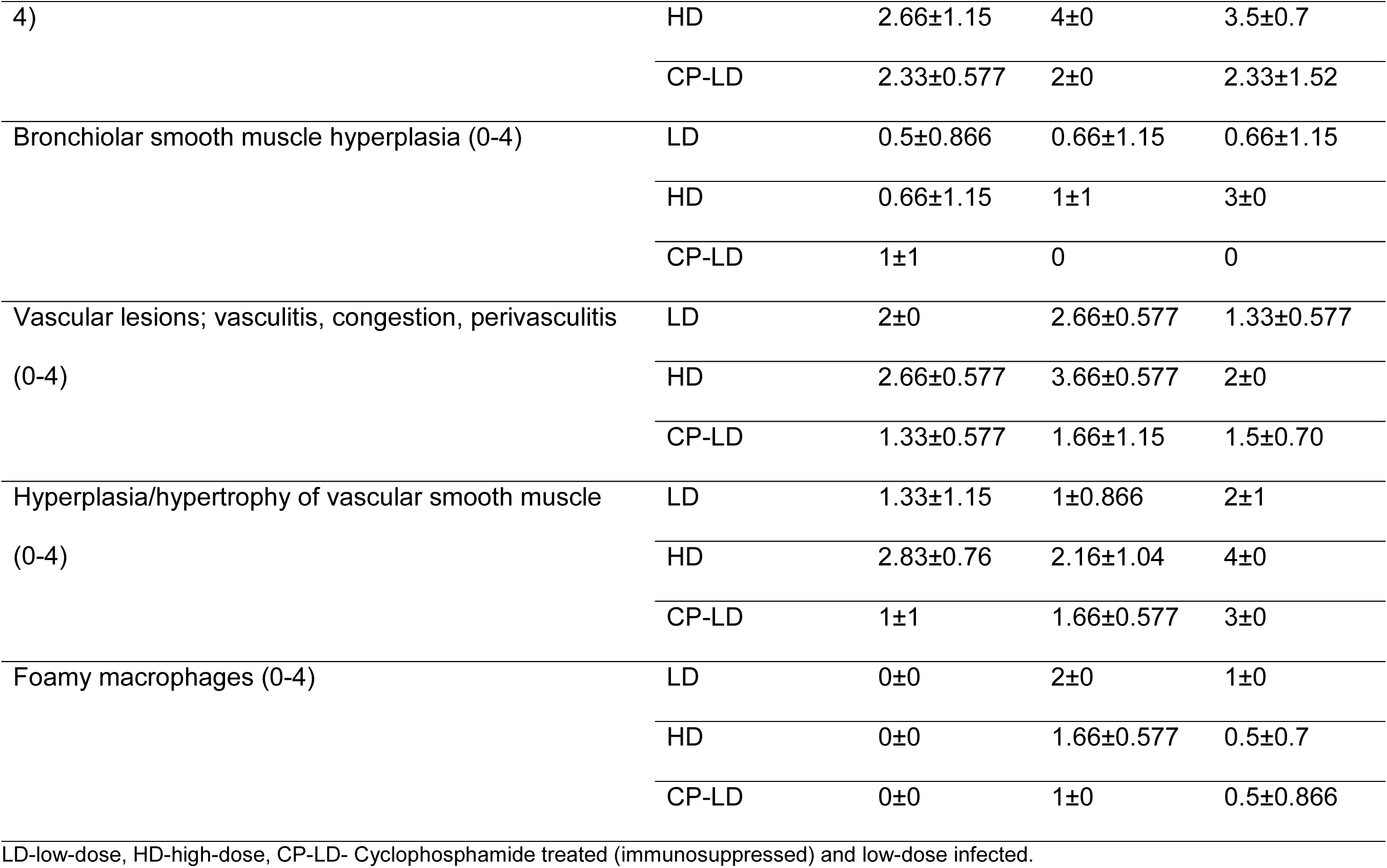
Histopathological scoring of SARS-CoV-2 infected hamster lungs.

The immunocompetent hamsters infected with HD SARS-CoV-2 had severe multifocal to diffused mononuclear cell infiltration in the interstitium at 4 dpi (Figure 2i, j, m, n). At this time, extensive bronchiolar hyperplasia with the detachment of hyperplastic epithelial cells and lymphocyte infiltration in the lumen (Figure 2i) and alveolar hyperplasia/ metaplasia, wherein alveolar cells were lined with cuboidal epithelial cells instead of the squamous epithelium was noted (Figure 2i, m). In addition, multifocal edema of the interstitium and alveoli, congestion of capillaries in the alveolar walls (Figure 2i, m), focal hemorrhages (Figure 2j, n), and foamy macrophages (Figure 2n) (1/3 hamsters) were observed (Figure 2b and Table 2). Trichrome staining of the lung sections showed multifocal thickening of alveolar walls. At 7dpi, severe pneumonia appeared as a pseudolymphoid tissue (Figure 2k, o), and infiltration of foamy macrophages (Figure 2o) in the pulmonary interstitial connective tissue was observed. At 7 and 16 dpi, lung sections showed “forming thrombi or early thrombi” in the arterioles (Figure 2g, h). However, a lesser degree of pulmonary interstitial pneumonia was noted at 16 dpi (Figure 2d, h, l, p) than at 7dpi (Figure 2c, g, k, o and Table 2). Overall, the degree of interstitial pneumonia and bronchiolar-alveolar hyperplasia was severe in HD SARS-CoV-2 infected hamsters compared to LD infected hamsters.

The immunocompromised SARS-CoV-2 infected hamsters (CP-LD) showed a lower number of inflammatory cells (Figure 2q-x) and edema in the interstitium (Figure 2q, u) than the immunocompetent/LD infected animals. In the CP-LD group, mild infiltration of foamy macrophages was observed at 7 dpi (Figure 2v). However, CP-LD hamsters revealed severe hyperplasia to metaplasia of bronchiolar-alveolar epithelial cells (Figure 2r, v, s, w) with occlusion of alveolar (Figure 2s, w) and bronchiolar lumen (Figure 2t, x) were observed throughout the lung parenchyma (Table 2).

The trachea of LD and HD infected immunocompetent hamsters revealed multifocal denudation of mucosal epithelial cells into the lumen, metaplasia of tracheal epithelium, hyperplastic goblet cells, infiltration of lymphocytes and neutrophils in the lumen and submucosa, mucous exudate in the tracheal lumen at 4 and 7 dpi (Supplementary Figure 1). On day 16, elevated mononuclear and neutrophil infiltration was noted in the LD group (Supplementary Figures 1a, d). However, the degree of neutrophil infiltration in the submucosa was higher, with some hamsters showing a complete detachment of mucosa-submucosal layer from the cartilage in the HD group (Supplementary Figures1b, e). In contrast, severe hyperplasia and necrosis of mucosal epithelia, severe infiltration of neutrophils, and mild lymphocytes infiltration in the submucosa were observed in the CP-LD hamsters as early as 4 dpi (Supplementary Figures1c, 1f). In these animals, multifocal necrosis and detachment of mucosal and submucosal layers were observed at 7 and 16 dpi (Supplementary Figures1c, 1f). Immunocompromised (CP-LD) hamsters had a lesser degree of inflammatory cells infiltration and the delayed resolution of inflammation in the lungs than LD-infected hamsters. Meanwhile, CP-LD hamsters showed severe bronchiolar-alveolar hyperplasia than LD-infected hamsters. It indicated that the disease is prolonged in CP-LD infected hamsters as observed in body weight change.

### 2.6. Hypertension-like vascular smooth muscle hyperplasia/hypertrophy in the lungs and kidney of SARS-CoV-2 infected hamsters

The arteriolar smooth muscle proliferation in the lungs and kidneys is associated with hypertension. We examined whether SARS-CoV-2 infected hamsters exhibit any of these pathological features. We observed thickening of the vascular smooth muscle layer/tunica media in the histology sections of the lungs (Figure 3a-f) and kidneys (Figure 3g-3l) of immunocompetent animals infected with LD (Figure 3a, 3d, 3g, 3j) and HD (Figure 3b, 3e, 3h, 3k) at 7 dpi. Two of the immunocompetent/HD-infected hamsters showed hypertrophy of the muscular layer in the arteriole and mononuclear cells infiltration in the adventitial layer of the arteriole (perivasculitis) and proliferation of tunica externa (Figure 3b, 3e). In contrast, in the immunocompromised LD infected hamsters, severe vascular lesions, denudation of endothelial cells into the lumen, mononuclear cells infiltration in the adventitia of small blood vessels, and rupture of the arteriolar wall, including muscular layer and leakage of vascular contents was noted at 7 dpi (Figure 3c, 3f).

**Figure 3.**
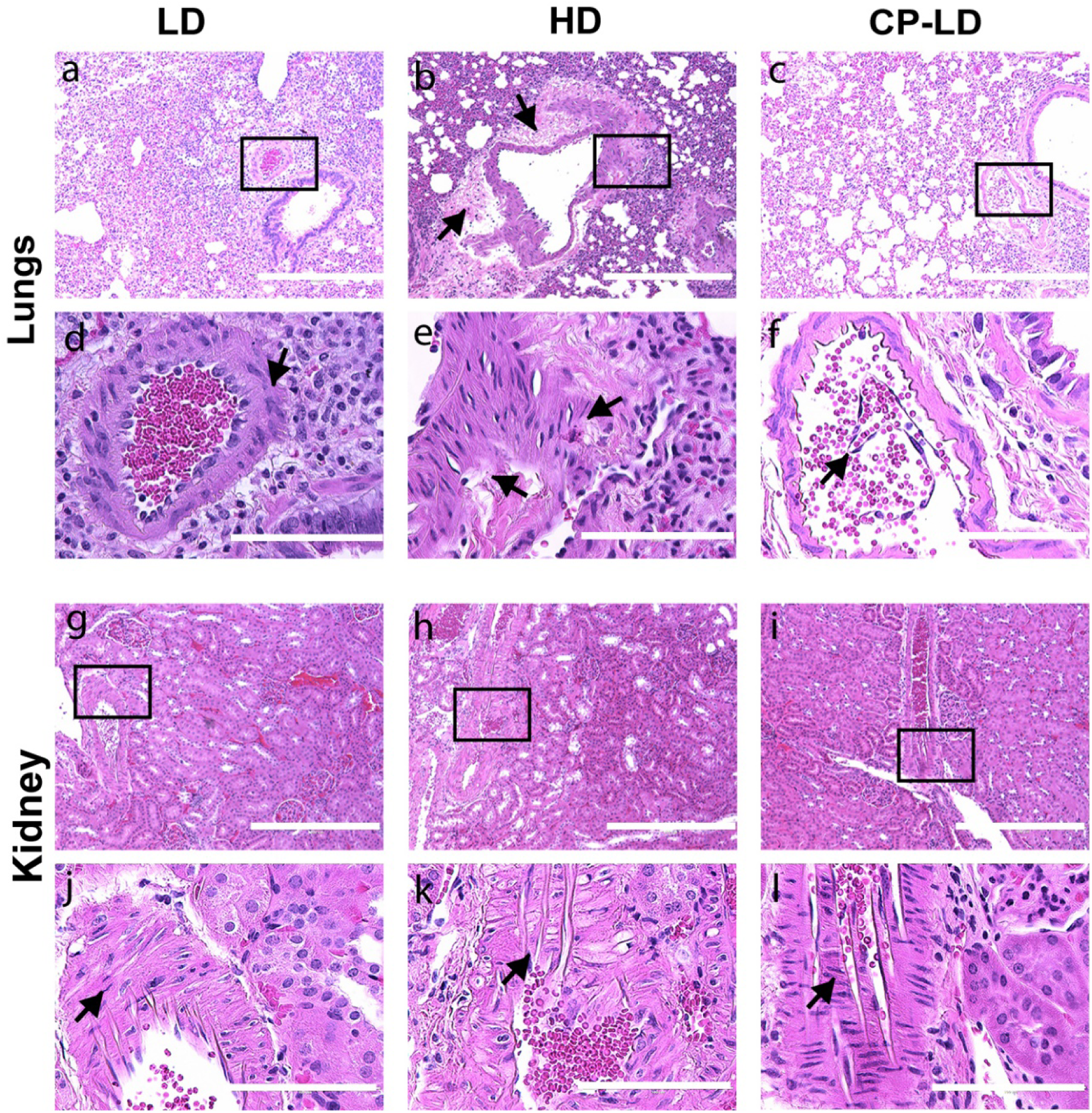
Arteriolar pathology in lungs and kidneys of SARS-CoV-2 infected hamsters. Histopathological analysis of pulmonary parenchyma revealed mild arteriolar smooth muscle hyperplasia **(a)**, severe perivasculitis and smooth muscle hyperplasia **(b;** *arrows*), detachment of endothelial cells into the arteriolar lumen, and mild perivasculitis **(c)** in LD, HD and CP-LD infected hamsters, respectively. The images **d**, **e**, **f** are a magnification of the boxed area in **a**, **b**, **c**, respectively. Moderate (LD and CP-LD) **(g**, **i)** to severe (HD) **(h)** vascular smooth muscle hyperplasia in the renal arterioles in SARS-CoV-2 infected hamsters. The images in **j**, **l**, **k** are a magnification of the boxed areas in **g**, **i**, **h,** respectively. The images **a-c**, **g-i** are 100x, and **d-f**, **j-l** are 400x magnifications (marked in 100x images). Scale bar represents 100 µm (**d-f**, **j-l**) or 400 µm (**a-c**, **g-i)**.

The LD, HD and CP-LD infected hamsters had severe adrenal cortical and medullary degeneration, necrosis and liquefaction (SupplementaryFigures2a-l) with cortical hypertrophy in some hamsters (Supplementary Figure 2e, f). A moderate-to-severe arteriolar smooth muscle hypertrophy and hyperplasia were also observed in the renal arterioles (Figure 3g-l). Thus, in addition to the hypertrophy/hyperplasia of vascular smooth muscles and adrenal cortical pathology, which are reported to be associated with hypertension, thickening of vascular smooth muscles can also be an indication of hypertension induced by SARS-CoV-2.

### 2.7. Multi-organ pathology induced by SARS-CoV-2 infection in immunocompetent and immunocompromised hamsters

Since steatosis (a.k.a. vacuolation) of hepatocytes is commonly observed among COVID-19 patients, we investigated these pathological features in the SARS-CoV-2infected hamsters ^21^. Liver histology revealed diffuse infiltration of a few lymphocytes and neutrophils in LD, HD and CP-LD infected hamsters (Figure 4a-4l) and degeneration of hepatocytes with pyknosis at 4 and 7 dpi (Figure 4). In addition, mild steatosis and portal vein congestion was observed at 16dpi (not shown). The LD-infected hamsters showed mild steatosis(Figure 4a, 4d); in contrast, immunocompetent HD SARS-CoV-2 infected hamsters showed moderate-to-severe multifocal to diffuse steatosis (Figure 4d) and portal vein congestion at 4, 7 and 16 dpi (Figure 4e, 4h).

**Figure 4.**
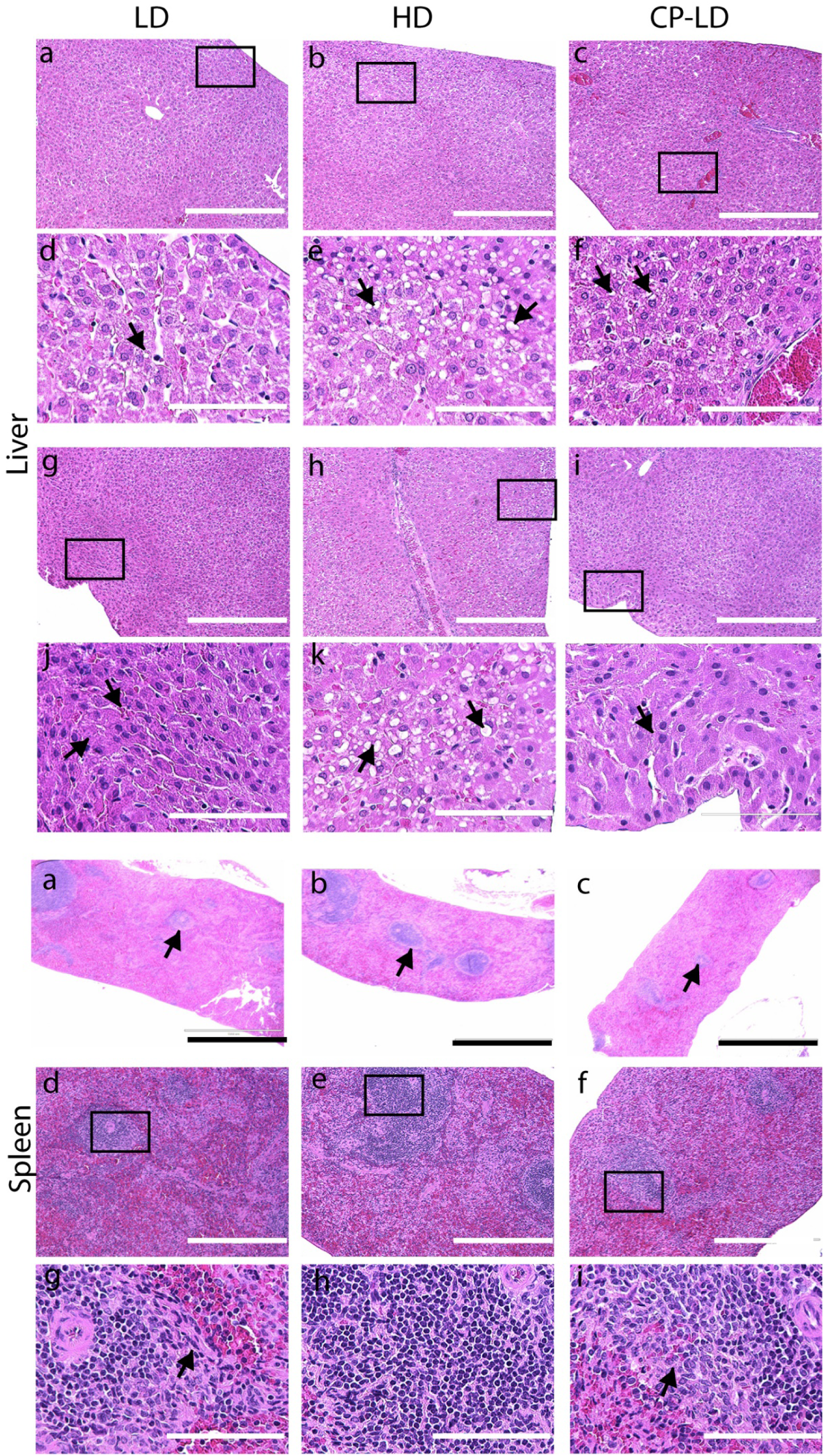
Histopathology of liver and spleen in SARS-CoV-2 infected hamsters. **A.** Histopathological analysis LD, HD and CP-LD infected hamster liver (4, 7 and 16 dpi, n=3 per time point) showed infiltration of lymphocytes in sinusoids at 4 dpi **(a-l),** and pyknosis (*arrows*), karyorrhexis, and karyolysis in necrotic hepatocytes **(g**, **j)** at 7dpi. Necrotic hepatocytes with pyknotic nuclei and mild to moderate steatosis (*arrows*) **(b**, **e**, **h**, **k)** and severe congestion in the portal vein **(h)** were noted at 4 dpi, while severe congestion in the portal vein, sinusoids, and mild steatosis **(c**, **f,** *arrows*) were observed at 7 dpi in HD infected hamsters. Severe degeneration and necrosis of hepatocytes near the parietal surface were observed at 4 dpi in CP-LD animals **(l,** *arrows*). Images **a-c**, **g-i** are 100x, and **d-f**, **j-l** are 400x magnifications. Scale bar represents 100 µm **(d-f, j-l)**, 400 µm **(a-c**, **g-i)**. Histopathological analysis of the spleen revealed a reduction in the number and area of white pulp lesions and cellular composition with increased trabecular connective tissues at 4 dpi in LD **(a**, **d**, **g)**, HD **(b**, **e**, **h)**, and CP-LD **(c**, **f**, **i)** infected hamsters. Images **a**, **b**, **c** are 40x, and **d**, **e**, **f** are 100x, and **g**, **h**, **i** are 400x magnifications. Scale bar represents 100 µm **(g**, **h**, **i)**, 400 µm **(d**, **e**, **f),** or 1000 µm **(a, b, c)**.

In contrast, in the CP-LD group, one of the three hamsters at 4 and 7 dpi, and all hamsters at 16 dpi had severe liver degeneration, marked with eosinophilic granular cytoplasm and pyknosis (Figure 4i, 4l). In these animals, mild-to-moderate steatosis, portal vein congestion, and edema in the sinusoids surrounding the portal vein were observed (Figure 4c, 4f).

Reduced size and cellular composition of splenic white pulp have been reported in COVID-19 patients ^22^. Therefore, we investigated this phenomenon in SARS-CoV-2 infected hamsters. Consistent with the clinical studies, the spleens of immunocompetent LD and HD SARS-CoV-2 infected hamsters showed a reduction in size and number of white pulp (Figure 4a-e, 4g, 4h) compared to uninfected hamsters at 4 dpi. In the infected animals, white pulp lymphocytes were replaced by trabecular connective tissues. In contrast, the immunocompromised LD SARS-CoV-2 infected hamsters (CP-LD) showed a high degree of white pulp atrophy at all the time points tested (4, 7 and 16 days post-infection) (Figure 4c, 4f, 4i).

The kidneys of immunocompetent hamsters infected with LD or HD showed mild lymphocyte infiltration in the interstitial tissues and multifocal tubular degeneration, acute tubular necrosis with the detachment of tubular cells into lumen, tubular epithelial cells degeneration with pyknotic nuclei or karyorrhexis or chromatolysis at 4, 7 and 16 dpi (Supplementary Figure3a-h). In these animals, focal regions of tubulointerstitial edema were also observed. The degree of acute tubular necrosis was higher in hamsters infected with HD (Supplementary Figure 3f) than LD (Supplementary Figure3d). At 7d pi (2 out of 3 hamsters) and 16 dpi (3 hamsters), a moderate renal arteriolar media (smooth muscle) hyperplasia/hypertrophy was observed in the LD-infected hamsters (Figure 3g, 3j). These pathological manifestations appeared early (4 dpi) in the HD-infected animals (Figure 3h, 3k).

The kidneys of CP-LD hamsters showed lymphocyte infiltration in the interstitial tissues, focal/multifocal acute tubular necrosis with the detachment of tubular epithelial cells from the basement membrane (Supplementary Figure 3i, 3j), forming concentric layers of eosinophilic masses with necrotic nuclei in the tubular lumen (Supplementary Figure 3k,3l), at 4, 7 and 16 dpi. However, the degree of tubular necrosis in these animals was less compared to the kidneys of immunocompetent LD-infected hamsters. Further, the arteriolar media hyperplasia/hypertrophy was less prominent in these animals (Figure 3i, 3l). The CP-LD hamsters also showed basophilic ground-glass bodies in the tubular epithelial cells and glomerular degeneration at 4, 7 and 16 dpi (Supplementary Figures3k, 3l).

### 2.8. SARS-CoV-2 host cell entry receptor expression in immunocompetent and immunocompromised hamster lungs

Since ACE-2 (angiotensin-converting enzyme-2) and CD147 are key receptors for SARS-CoV-2 host cell entry, we investigated the expression of ACE2, CD147 as well as SARS-CoV-2 in the lung cells using smRNA-FISH analysis (Figure 5). We observed a slightly elevated level of SARS-CoV-2 N protein in the immunocompromised compared to immunocompetent infected hamsters at 4 (Figure 5c, d) and 16 dpi (Figure 5e, f), though the difference in the number of cells positive for the SARS-CoV-2 N gene was not statistically significant (Figure 5s). At both 4 and 16 dpi, ACE2 receptor expression was insignificantly lower in the infected immunocompetent and immunocompromised hamsters compared to the respective uninfected control groups (Figure 5g-l, 5t). In contrast, CD147 expression was higher in the infected immunocompromised hamsters at both 4 and 16 dpi, compared to the infected and uninfected immunocompetent animals. However, the difference was not statistically significant between these groups (Figure 5m-r, 5u). Together, the data suggest that key SARS-CoV-2 host cell entry receptors (ACE-2 and CD147) show distinct expression patterns between immunocompetent and immunocompromised hamster lungs, although the difference in expression pattern did not correlate with the disease pathology in respective infection groups.

**Figure 5.**
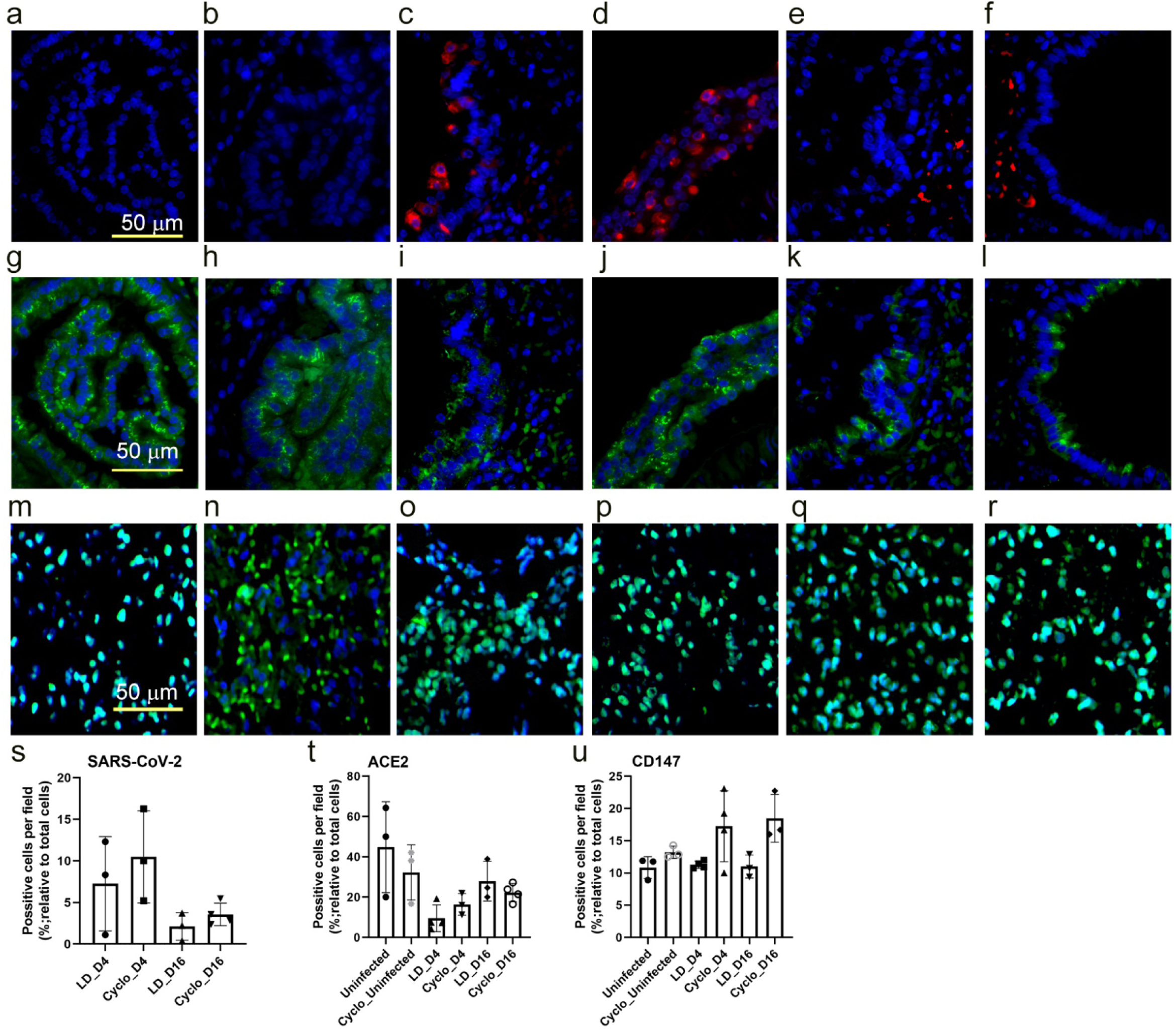
Spatial expression of SARS-CoV-2 host cell entry receptors in immunocompetent and immunocompromised hamsters. Lung sections were stained for SARS-CoV-2 **(a-f)** or ACE-2 receptor **(g-l)** or CD147 **(m-r)**. Images **a, g, m** are uninfected immunocompetent hamster lung sections, and **b, h, n** are uninfected immunocompromised hamster lung sections. Images **c, i, o** are SARS-CoV-2 infected immunocompetent hamster lung sections at 4 dpi; and **d, j, p** are SARS-CoV-2 infected immunocompromised hamster lung sections at 4 dpi. Images **e, k, q** are SARS-CoV-2 infected immunocompetent hamster lung sections at 16 dpi; and **f, l,r** are SARS-CoV-2 infected immunocompromised hamster lung sections at 16 dpi. The red color in **c-f** indicates the presence of SARS-CoV-2, and the green color indicates ACE-2 receptor expression in **g-l**, or CD147 expression in **m-r**. Image s shows lung cells positive for SARS-CoV-2 in immunocompetent (LD) and immunocompromised (Cyclo) hamsters at 4 dpi (D4) and 16 dpi (D16); image t shows lung cells positive for ACE-2 in uninfected and SARS-CoV-2 infected immunocompetent (LD) and immunocompromised (Cyclo) hamsters at 4 dpi (D4) and 16 dpi (D16) and image u shows lung cells positive for CD147 in uninfected and SARS-CoV-2 infected immunocompetent (LD) and immunocompromised (Cyclo) hamsters at 4 dpi (D4) and 16 dpi (D16). n=3 per group.

### 2.9. SARS-CoV-2 infection elicits distinct transcriptome profiles in the lungs

To gain insight into the pulmonary response to SARS-CoV-2 infection, we performed RNAseq analysis of immunocompetent and immunosuppressed hamster lungs at 4 dpi (acute) and 16 dpi (chronic). The data from infected animals were normalized to the respective group of uninfected animals (i.e., with or without immunosuppression). The principal component analysis (PCA) showed segregation of uninfected groups from those at different stages of infection with or without immunosuppression (Figure6a, b). Analysis of differentially expressed genes showed a greater number of significantly differentially expressed genes (SDEG) at 4dpi than 16dpi in both immunocompetent and immunocompromised animals upon SARS-CoV-2 infection (Figure6c). However, the number of SDEGs in the immunosuppressed animals was less than in immunocompetent animals at 4dpi (1,273 versus 1,829). In contrast, the former group had more SDEGs at 16dpi (646 versus 273). Thus, immunocompetent and immunocompromised hosts showed distinct gene expression profiles at different stages of infection (i.e. 4dpi/acute and 16dpi/chronic stage). Among the 4 groups (i.e.4dpi and 16dpi in immunocompromised and immunocompetent groups), 42 common SDEGs were identified. Of these, 31 were upregulated in the immunocompetent group, compared to 10 SDEGs in the immunocompromised group at 4 and 16 dpi, respectively(Figure6d). Furthermore, 50% of the 42 SDEGs were expressed in opposite directions between the immunocompetent and immunocompromised groups at 4 and 16 dpi.

**Figure 6.**
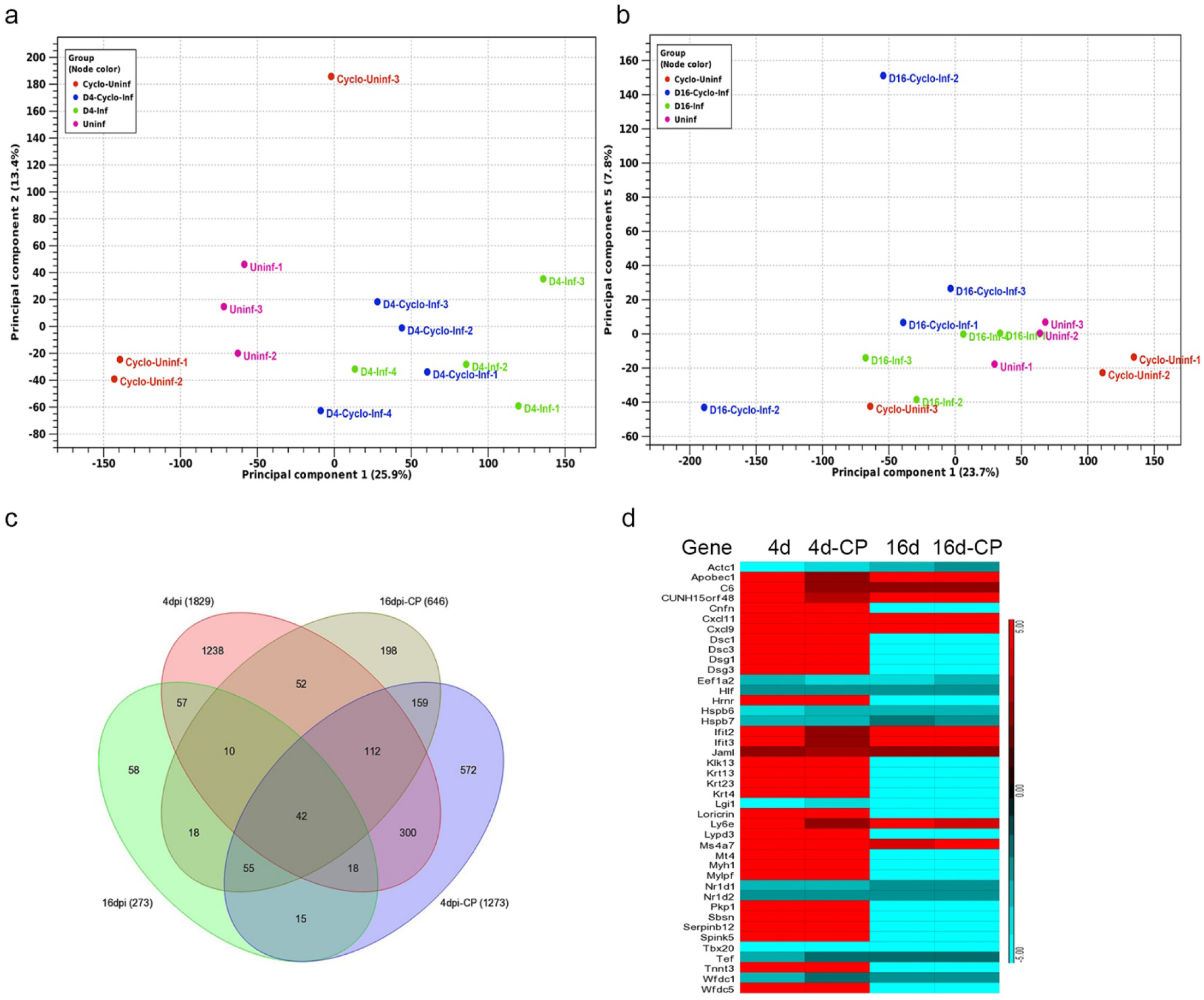
Genomewide lung transcriptome analysis of immunocompetent and immunocompromised hamsters infected with SARS-CoV-2. **a.** PCA plot of immunocompetent and immunocompromised hamsters without infection or 4 dpi. **b.** PCA plot of immunocompetent and immunocompromised hamsters without infection or 16 days post SARS-CoV-2 infection. **c.** Venn diagram showing the number of significantly differentially expressed genes (SDEG) in the lungs of immunocompetent and immunocompromised hamsters at 4 dpi or 16 dpi. Data from the infected animals were normalized to corresponding uninfected animal data. **d**. Heat-map of SDEGs commonly perturbed in all four groups. Scale bar shows up (red) and down-regulation (cyan). n=3 per group.

### 2.10. SARS-CoV-2 infection elicits distinct acute and chronic transcriptome profiles in the lungs of the immunocompetent host

Next, we analyzed the gene networks and pathways differentially affected by SARS-CoV-2 infection at 4dpi and 16 dpi in immunocompetent animals. Canonical pathways upregulated at 4 dpi strongly suggests the induction of a classical innate and proinflammatory host response. This includes natural killer (NK) cell signaling, hyper cytokinemia/chemokinemia, pattern recognition receptor (PRR) signaling, dendritic cell (DC) maturation, communication between DC and NK cells, and Th1 pathway (Supplementary Figure 4). Interferon (IFN) lambda-3 and IFN beta-1 were among the top upregulated genes in this group (Figure 7). In contrast, the top canonical pathways perturbed at 16 dpi involved tissue thrombosis, such as intrinsic prothrombin activation, MSP-RON signaling in macrophages and coagulation system pathways (Supplementary Figure 5). Other pathways involved in host cell activation, including LXR/RXR activation, production of reactive oxygen and nitrogen intermediates in macrophages, and acute phase response signaling, were significantly dampened at this time when the animals recovered from viral infection (i.e, no live virus) and gained body weight. Thus, the genomewide lung transcriptome profile is consistent with the corresponding pathophysiological manifestations observed during acute (4dpi) and chronic (16 dpi) stages of infection. Notably, expression of the SDEGs involved in hypercytokinemia/chemokinemia (e.g., *CCL5, IFNB1*, *IFNG, IFNL3, IL1B, IL1, IL18, IRF7* and *IRF9*) and interferon signaling pathways (e.g., *CXCL9*, *CXCL11*, *IFIT2*, *IFIT3* and *APOBEC1*) were upregulated at 4 and 16 dpi in infected immunocompetent hamsters (Figure7e, f and Supplementary Figures 6 and 7).

**Figure 7.**
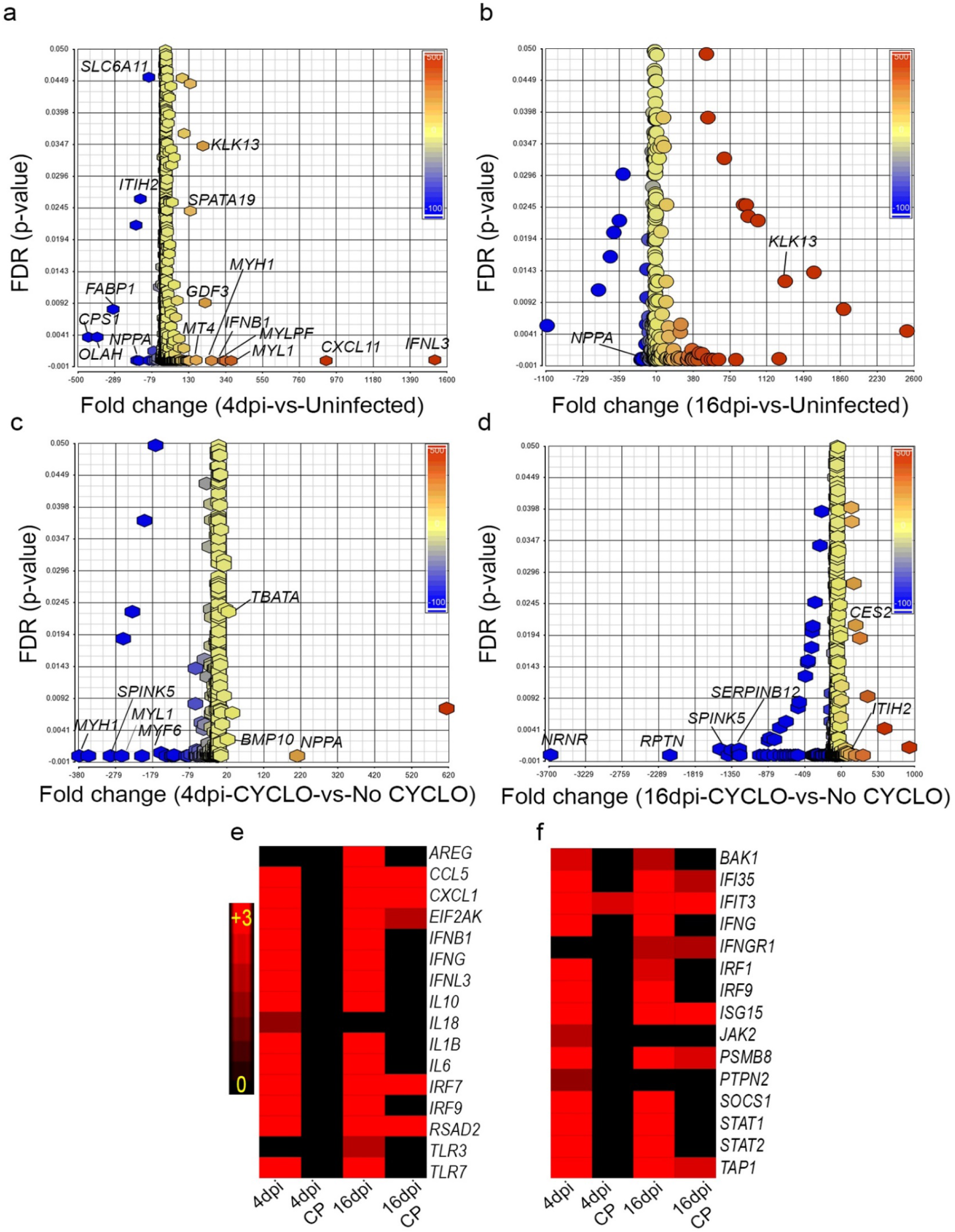
Distribution of differentially expressed genes in immunocompetent and immunocompromised hamsters infected with SARS-CoV-2. **a.** Scatter plot of SDEG in immunocompetent hamster lungs at 4 dpi. **b.** Scatter plot of SDEG in immunocompetent hamster lungs at 16 dpi **c.** Scatter plot of SDEG in immunocompromised hamster lungs at 4 dpi. **d.** Scatter plot of SDEG in immunocompromised hamster lungs at 16 dpi. **e.** Heat-map of SDEG involved in hypercytokinemia/hyperchemokinemia network. f. Heat-map of SDEG involved in canonical IFN signaling pathway. Scale bar shows up-regulation (red) and no expression (black). FDR-false discovery rate.

Transcriptomes inSARS-CoV-2 infected immunocompetent hamsters and uninfected controls at 4 dpi were subjected to Ingenuity Pathway Analysis (IPA). Biological processes affected included upregulation of host cell death, cytotoxicity, antimicrobial response, various types of immune cell development, recruitment and activation, and the inflammatory response. In contrast, biological functions associated with organismal survival were dampened in these animals (Supplementary Table1). At 16 dpi, the biological processes related to immune cell infiltration, edema, tissue necrosis, and organismal death were upregulated. At the same time, other functions related to lipid metabolism, including fatty acid metabolism and lipid transport, as well as the survival of organisms, were dampened in the infected immunocompetent hamsters, compared to uninfected controls (Supplementary Table2). In contrast, analysis of immunocompromised hamster lung transcriptomes indicated that host biological functions associated with organismal survival, cell signaling, molecular transport, cell-mediated immunity, immune cell trafficking, and function were dampened, while cell death-related functions, such as apoptosis, were upregulated at 4 dpi (Supplementary Table3). However, some of these functions, including cell movement and molecular transport, and cell function and maintenance, were upregulated in the infected immunocompromised hamster lungs at 16 dpi. Importantly, humoral immune response functions, such as the number of B lymphocytes and quantity of immunoglobulin (Ig) was significantly downregulated in these animals (Supplementary Table4). This observation is consistent with the loss of antibody response in the infected immunocompromised animals. Together, the biological functions that are significantly perturbed in the immunocompetent and immunocompromised hamster lungs upon SARS-CoV-2 infection are consistent with and support the disease pathology and immune/antibody response in the respective animals.

### 2.11. Dampened proinflammatory response pathways in the lungs of SARS-CoV-2 infected immunocompromised hamsters

To determine the differential regulation of gene networks and pathways in the lungs of immunocompromised versus immunocompetent hamsters infected with SARS-CoV-2, we interrogated the RNAseq data at 4 dpi and 16 dpi between these two groups after normalization to uninfected controls. Dampening of proinflammatory, innate immune function pathways, including phagosome formation, production of reactive oxygen and nitrogen species (ROS and RNS), dendritic cell maturation and NK cell activation was noted at 4 dpi in the immunocompromised, compared to immunocompetent animals (Figure 8a, b and c, and Supplementary Figure 8). At 16 dpi, network genes involved in MPN-RON signaling and prothrombin signaling were upregulated in the SARS-CoV-2-infected immunocompetent hamsters (Figure 8d, e and Supplementary Figure 9). In contrast, LXR/RXR signaling was upregulated in the infected immunocompromised hamsters (Figure 8f and Supplementary Figure 9). However, the B cell receptor signaling and the network genes that determine the number of B cells were downregulated in this group (Figure 8g). This observation is consistent with and supported by the lack of antibodies in the sera of infected immunocompromised hamsters.

**Figure 8.**
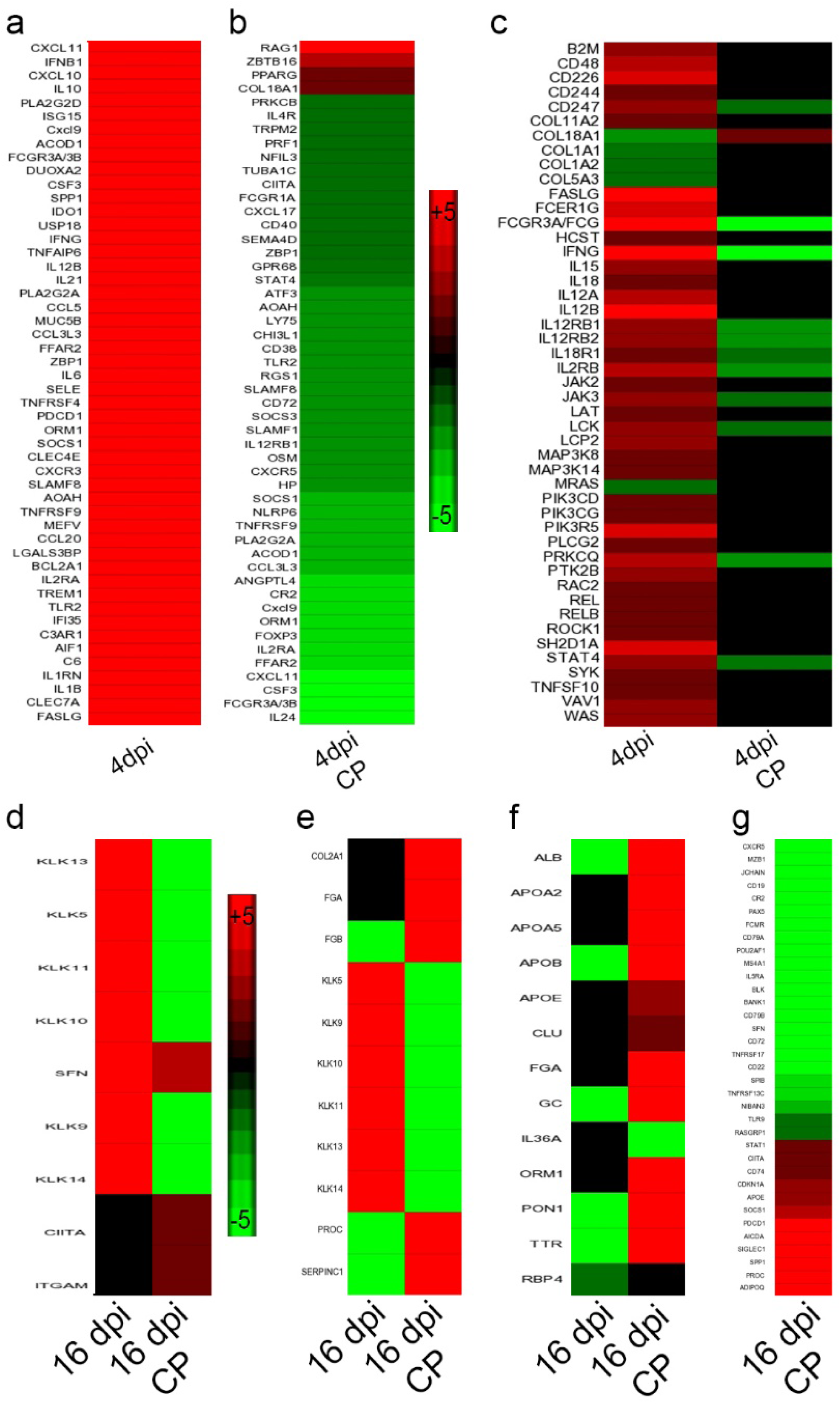
Expression of network genes in immunocompetent and immunocompromised hamsters infected with SARS-CoV-2. **a.** Heat map of SDEG involved in the inflammatory response in immunocompetent hamster lungs at 4 dpi. **b.** Heat map of SDEG involved in the inflammatory response in immunocompromised hamster lungs at 4 dpi (4 dpi CY). **c.** Heat map of SDEG involved in NK cell activation network in immunocompetent (4 dpi) and immunocompromised (4 dpi CY) hamster lungs at 4 dpi. **d.** Heat map of SDEG involved in canonical MPN-RON signaling in macrophages network in immunocompetent (16 dpi) and immunocompromised (16 dpi CY) hamster lungs at 16 dpi. **e.** Heat map of SDEG involved in prothrombin signaling network in immunocompetent (16 dpi) and immunocompromised (16 dpi CY) hamster lungs at 16 dpi. **f.** Heat map of SDEG involved in canonical LXR/RXR signaling pathway in immunocompetent (16 dpi) and immunocompromised (16 dpi CY) hamster lungs at 16 dpi. **g.** Heat map of SDEG involved in B cell recruitment and accumulation network in immunocompetent (16 dpi) and immunocompromised (16 dpi CY) hamster lungs at 16 dpi. Scale bar shows up (red) and down-regulation (green).

## 3. Discussion

The differential pathogenesis of COVID-19 between immunocompromised and immunocompetent individuals is poorly understood^23^. Similarly, predictors of disease severity, including the pathological manifestations of extrapulmonary organs upon SARS-CoV-2 infection in immunocompetent and immunocompromised hosts, are not well described. The Syrian hamster model has been an invaluable tool to understand the host-SARS-CoV-2 interactions and disease pathogenesis as well as to devise intervention strategies to combat COVID-19 ^14–17^. Here, we report the dynamic changes in SARS-CoV-2 replication and pathological manifestations in pulmonary and extrapulmonary organs of immunocompetent and immunocompromised hamsters. We also report the molecular correlates of the host response to infection with different initial inoculum doses of SARS-CoV-2 in immunocompetent hamsters. Together, our data suggest that the histopathological manifestations caused by progressive SARS-CoV-2 infection predict COVID-19 severity better than individual measures of viral load, antibody response, and cytokine storm at the systemic or local (lungs) levels in the immunocompetent and immunocompromised hosts.

In patients with COVID-19, primarily a respiratory disease, the highest viral load has been reported in the lungs. However, low levels of viral RNA were detected in the cardiovascular, endocrine, gastrointestinal, urogenital, hemopoietic, and central nervous systems ^24, 25^. In most COVID-19 patients, the infectious viruses were not isolatable after 10 days of the onset of symptoms despite detectable viral RNA^26^. Consistent with these reports, we observed the highest viral load in the pulmonary system and detected extrapulmonary dissemination of SARS-CoV-2 to other organs. Notably, a similar viral load was observed in various LD and HD infected hamster tissues, despite a 5 log difference in the initial infectious inoculum between these two doses. In these animals, infectious viruses were observed in lungs up to 7 dpi; however, the SARS-CoV-2 N gene transcript was detected until 16 dpi. These findings were supported by a previous study on the hamster model by Imai et al.^14^. Together, these observation suggests that the kinetics of viral replication and persistence in hamsters recapitulates the findings in COVID-19 cases.

In humans, the neutralizing antibodies generated following exposure to SARS-CoV-2 or vaccination constitute a significant determinant of virus clearance and protection^13, 27^. Most of the SARS-CoV-2 infected individuals develop antiviral antibodies by 7-14 days ^28^. In addition, clinical data shows that the high viral load and the severe disease correlate with increased antibody titer among COVID-19 cases ^27, 29^. Consistently, in our hamster studies, neutralizing antibodies were present at 7 dpi, and virus clearance coincided with the appearance of these serum neutralizing antibodies. The virus neutralizing antibody titer in these hamsters was proportional to the initial infectious dose of the virus, as reported in other studies ^15, 30^.

Patients with eosinophilic granulomatosis and polyangiitis are treated with cyclophosphamide therapy. These patients did not produce any SARS-CoV-2 specific antibodies after they acquired symptomatic COVID-19^31^. Likewise, we did not observe any SARS-CoV-2 neutralizing antibodies in the sera of infected immunocompromised hamsters, which was also reported in a previous study ^32^. Consequently, no virus clearance was observed in the SARS-CoV-2 infected immunocompromised animals up to 16 dpi. In this study, lung RNA expression analysis on 16 dpi revealed that the B cell activation network genes, including CD19, CD22, CD72, FcgR, were downregulated among the immunocompromised hamsters infected with SARS-CoV-2. It was reported that the cyclophosphamide treatment could inhibit B cell activation, proliferation, differentiation, and immunoglobulin secretion in patients undergoing cyclophosphamide therapy ^33, 34^. The immunosuppression treatment of hamsters with cyclophosphamide treatment would likely have diminished the number of T and B cells and/or abolished the ability to produce antibodies upon antigen exposure^31^.

Consistent with this notion, we observed dampened IL4 expression, essential for B cell differentiation, in the infected immunocompromised animals. Furthermore, severely atrophied splenic lymphoid follicles were noted in the immunocompromised, compared to immunocompetent hamsters. Thus, the lack of IL4-mediated B and/or T cell activation, and therefore the antibody production during SARS-CoV-2 infection, might have compromised the onset of an effective antiviral response in the immunosuppressed host. Similarly, the SARS-CoV-2 infected immunocompromised hamsters had mild pulmonary lymphocytic infiltration and poor resolution of pneumonia and sustained a prolonged weight loss, compared to immunocompetent hamsters, which is consistent with previous observations in hamsters^14, 32^. The elevated IL-10 expression in the immunocompromised hamsters might have played a role in reducing the immune cell infiltration to the site of infection (i.e., the lungs).

A pathologic hallmark of severe COVID-19 cases is the onset of inflammatory “cytokine storm”, marked by elevated IL1B, TNFA, CCL2, IL6, MIP1A, and IL10 in the plasma ^35^. Severe COVID-19 was also correlated with bodyweight loss ^36^. Consistent with these reports, we observed elevated inflammatory cytokine levels in the infected immunocompetent and immunocompromised hamsters. Furthermore, bodyweight loss was significantly higher among hamsters infected with a high dose of the virus. These animals also showed more robust inflammation and disease pathology than low-dose infected animals. Although, the HD-infected hamsters showed more severe hyperinflammation than LD, the expression cytokines in LD and HD were not significantly different except IFNG (4 dpi) and IL4 (16 dpi). Thus, it appears that once active disease is established at about 4 dpi, the expression pattern of many of the cytokine storm molecules does not correlate with the infectious dose, or the degree of pathological manifestations in the SARS-CoV-2 infected hamsters.

The lungs of immunocompetent hamsters infected with SARS-CoV-2 revealed upregulation of innate immune and inflammatory pathway genes, including IFNB and IFNL, NK cell activation, hypercytokinemia as observed in previous studies ^20^. However, the immunocompromised SARS-CoV-2 infected hamsters showed a dampening of innate immune and inflammatory responses, including NK cell activation. The poor innate immune response correlated with virus persistence in immunocompromised hamsters. Lower levels of inflammation in the immunocompromised hamsters could be due to the lack of optimal activation of proinflammatory pathways. Cyclophosphamide treatment has been shown to inhibit Treg cell functions^37^, and low dose cyclophosphamide treatment in mice was reported to cause reduction and alteration in immune cells in spleen and lymph nodes ^20, 38^. The hyper induction of proinflammatory cytokines such as IL6, IL1B, IFNG was observed in COVID-19 patients, and the level of induction was proportional to the severity of disease ^35^. However, the causal link between cyclophosphamide treatment and reduced proinflammatory cytokines and inflammation levels in COVID-19 cases remains unknown. Since the treatment regimen for COVID-19 patients includes a combination of anti-inflammatory agents, it is challenging to determine the differential immunomodulatory effect due to the treatment versus SARS-CoV-2 infection. The hamster model will allow these effects to be teased apart in future work.

Clinical studies show that a significant proportion of COVID-19 patients had bronchial/bronchiolar wall thickening in computed tomography (CT) scan reports ^39, 40^. In general, the thickening of bronchiolar smooth muscles is a pathognomonic characteristic of asthma, which is mediated by a TH2 response, marked with elevated levels of IL4 and airway inflammation ^41^. Further, CT scans from COVID-19 patients showed thickening of blood vessels in lungs, rupture of pararenal aortic aneurism, and cerebral aneurysm^42, 43^. Although rhinoviruses have been reported to exacerbate asthma in infected patients ^44, 45^, whether asthma is induced/exacerbated in COVID-19 patients remains unclear. We observed the thickening of bronchiolar smooth muscles in SARS-CoV-2 infected hamsters in an infectious-inoculum dose-dependent manner. In addition, as reported in human clinical studies^46^, we observed extensive smooth muscle hypertrophy/hyperplasia in the SARS-CoV-2 infected hamster pulmonary arterioles and renal arterioles and rupture of pulmonary vessels in some animals. While the exact mechanism underlying vascular thickening during SARS-CoV-2 infection is not fully understood, disruption of the renin-angiotensin system during COVID-19 could contribute to this anomaly ^47^.

Autopsies have revealed that cortical degeneration and necrosis, adrenalitis, and cortical hyperplasia of adrenal glands are associated with COVID-19 infection ^48, 49^. Similar adrenal cortical and medullary lesions were also observed in the hamsters, affecting deregulating blood pressure during SARS-CoV-2 infection ^50, 51^. However, the causal association between adrenal cortical/medullary insufficiency and the severity of COVID-19 is yet to be unraveled. Clinical studies have also shown microvesicular hepatic steatosis among COVID-19 cases with or without underlying health conditions ^52, 53^. Although previous studies in the hamster model did not report any abnormality in the liver ^16, 17^, we observed mild degeneration and steatosis in both LD and HD SARS-CoV-2- infected hamsters. The discrepancy between these studies could be due to the inherent differences in the experimental design and the extent of pathological analysis performed.

Severe COVID-19 is associated with acute kidney injury and renal failure ^54–56^. Autopsy reports indicate the presence of proximal convoluted tubular epithelial necrosis, loss of brush border, detachment into the lumen of the kidney in patients who died of COVID-19 ^54^ and during biopsy ^55^. Additional symptoms of acute kidney injury, including proteinuria and elevated serum creatinine levels, were also noticed in these cases ^55, 56^. Consistent with these reports, we observed acute tubular epithelial necrosis of kidneys following SARS-CoV-2 infection in immunocompetent and immunocompromised hamsters. However, the precise mechanism of renal injury during COVID-19 is yet to be determined.

In summary, our findings reveal the kinetics of viral replication, antibody response, and associated disease pathology of various internal organs in immunocompetent and immunocompromised hamsters following SARS-CoV-2 infection. The histopathologic findings in our hamster models closely mimic the clinical and pathological manifestations observed in human COVID-19 cases. The two hamster models described in this work could be used to unravel the pathogenesis, tissue injury, viral transmission within and outside of the infected host. These models can also serve as a preclinical tool to evaluate potential intervention strategies such as therapeutics and vaccines to combat the ongoing COVID-19 pandemic.

## 4. METHODS

### 4.1. Virus and cell lines

SARS-CoV-2 (strain USA-WA1/2020) infected Vero E6 cell lysate was obtained from BEI Resources (BEI Resources, Manassas, VA, USA). Virus propagation, virus titration, infectivity assays, and antibody titration were performed using Vero E6 cells (ATCC, Manassas, MA, USA) ^57, 58^. Experiments involving infectious SARS-CoV-2 were conducted in Biosafety level 3 facilities at Rutgers University as per approved standard operating procedures. Unless specified, all chemicals and reagents were purchased from Sigma-Aldrich (Sigma-Aldrich, St. Louis, MO, USA).

### 4.2. Virus propagation and infectivity titration

We performed a viral plaque-forming unit (PFU) assay to determine live viruses in the inoculum and tissue homogenates. Vero E6 cell monolayer was infected with SARS-CoV-2 at a multiplicity of infection (MOI) of 0.2 in DMEM media (supplemented with 2% FBS). After 48 hours of infection, the supernatant from infected cells was collected and stored at -80 °C. For the PFU assay, Vero E6 cells were seeded onto 6-well plates at 5×10^5^ cells per well for 18-24 hours. The next day, an aliquot of virus supernatant was thawed at 37 °C and made 10-fold dilutions from 10^-2^ to 10^-6^. The spent media from 6-well plates were aspirated and infected with each virus dilutions as 400µL/well at 37°C for 1 hour. Unattached viruses were then removed, and the infected monolayers were overlaid with a mixture containing equal amounts of 2X MEM with 8% FBS and 1.6% low-melting agarose. The plaques were visualized on the 3rd day by staining with 0.2% crystal violet.

### 4.3. Golden Syrian hamster infection and sample collection

Seventy-five (n=75) male Golden Syrian hamsters (*Mesocricetusauratus*) between 6-8 weeks old were purchased (Envigo corporation, Denver, PA, USA) and housed at two animals/cage. Feed and water were given adlibitum throughout the experiment. Animals were acclimatized for seven days in the BSL3 facilities. Bodyweight, food and water intake were monitored twice a day for each animal throughout the experiment.

#### Immunocompetent hamsters

For intranasal infection, SARS-CoV-2 were prepared in two doses; low dose (LD;10^2.5^ PFU) and high dose (HD;10^6^ PFU) in 50µL of sterile 1X PBS. LD was administered to 30 animals, and HD was inoculated to 15 healthy hamsters. For the uninfected control group, five hamsters were intranasally inoculated with 50µL of sterile 1xPBS.

#### Immunosuppression treatment of hamsters

Cyclophosphamide was injected intra-peritoneally into 30 animals at 70mg/kg on the day of intranasal SARS-CoV-2 inoculation with 10^2.5^ PFU (CP-LD) and then every three days until the end of the experimental time point (16 dpi). This method has previously been shown to immunocompromise hamsters and causes them to be more vulnerable to progressive SARS-CoV infection ^34^. Five animals treated with cyclophosphamide, as mentioned above, were intranasally inoculated with 50µL of sterile 1xPBS as the control group.

Six animals from LD and cyclophosphamide-LD (n=6) groups and three animals from the HD group were euthanized on 2, 4, 7, 12 and 16 days post-infection. Blood was collected by cardiac puncture before autopsy. The turbinates, larynx and trachea, lung, heart, liver, spleen, adrenal, kidney, colon, epididymal fat, brain, eyes were collected and weighed aseptically. A portion of the harvested tissues was transferred to DMEM containing penicillin-streptomycin and was homogenized in a FastPrep bead beater (MP Biomedicals, Solon, OH, USA) as five cycles of 20 seconds each. A portion of the lung, trachea, liver, spleen, kidney, intestine, pulmonary artery, a hemisphere of the brain, one testis, eye, and adrenal gland were collected in 10% buffered formalin for histopathology analysis. A portion of the lung was stored in Trizol (ThermoFisher Scientific, CA, USA) at -80 °C for RNA extraction.

All animal procedures were performed in bio-safety level 3 facilities following procedures approved by the Rutgers University Institutional Animal Care and Use Committee, which is consistent with the policies of the American Veterinary Medical Association (AVMA), the Center for Disease Control (CDC) and the United States Department of Agriculture (USDA).

### 4.4. Histopathology analysis

The tissues were fixed in 10% buffered formalin, made into paraffin blocks, sectioned to 5-micron thickness, and stained with hematoxylin and eosin or Trichrome as described previously ^59^. The identity of samples was blinded before analysis, and histopathological examination was performed by a veterinary pathologist (S.R) using the EVOS FL Cell imaging system (Thermo Fischer Scientific, USA). Histopathology images were organized and labeled using Adobe Photoshop v22.1.1 and Adobe Illustrator v25.1. Spleen white pulp cells were quantified manually and using ImageJ (NIH, Bethesda, USA). The pulmonary pathology was scored (0-4/5) based on the degree of mononuclear infiltration, edema, alveolar/bronchiolar hyperplasia and metaplasia, emphysema, vascular lesions, bronchiolar/arteriolar smooth muscle thickening, and foamy macrophages. The lowest and highest score indicates mild to severe lesions (Table 2).

### 4.5. Immunohistochemistry

The formalin-fixed, paraffin-embedded tissue blocks were sliced into 5-micron sections and processed following standard procedures^59^. Briefly, the sections were immersed in xylene and descending grades of ethanol, and antigen retrieval was performed using 10mM citrate buffer at 90°C for 40 minutes. The sections were blocked using 1% BSA for 1 hr and incubated with rabbit anti-hamster ACE2 antibody (Product no. HPA000288, Millipore Sigma, Burlington, MA, USA) overnight at 4°C. Alexa-488 labeled anti-rabbit secondary antibodies (Cat no. ab150077, Abcam, MA, USA) prepared at 1:2000 dilution in 5% BSA were used to detect ACE2 expression. The slides were washed with 2X SSC buffer containing 10% formamide in 2X SSC (ThermoFisher Scientific, Waltham, MA, USA), followed by hybridization buffer containing 10% dextran sulphate, 1mg/mL *E.coli*tRNA, 2 mM ribonucleosidevanadyl complexes (New England Biolab, Ipswich, MA, USA), 0.02% ribonuclease free bovine serum albumin (ThermoFisher Scientific, Waltham, MA, USA), 10% formamide, 2X SSC for 30 minutes. Labeled smRNA-FISH probes (Biosearch Technologies, Dexter, MI, USA) were added and incubated overnight in a moist chamber at 37°C. The slides were washed in wash buffer, treated with TrueBlack Lipofuschin autofluorescence quencher (BiotiumInc, Fremont, CA, USA), and mounted with a coverslip. Images were captured using Axiovert 200M inverted fluorescence microscope (Zeiss, Oberkochen, Germany) using 20X or 63X oil-immersion objective with Prime sCMOS camera (Photometrics, Tucson, AZ) by Metamorph image acquisition software (Molecular Devices, San Jose, CA) ImageJ software was used for analyses. The number of cells positive for a marker was normalized to the total number of cells in each field (3-5 fields containing at least 500 cells were analyzed per sample). Further, GraphPad Prism-8 (GraphPad Software, San Diego, CA) was used for the statistical analysis of data. P values <0.05 were considered statistically significant.

### 4.6. Virus infectivity assays on tissues

Tissue homogenates were centrifuged, and the supernatant was filtered through a 0.45 µ filter. The filtrate was diluted in serum-free DMEM, and 400 µL was used to infect the one-day-old Vero E6 cell monolayers in the six-well plates as mentioned above.

### 4.7. Plaque reduction neutralization (PRNT) assay

Hamster serum samples were heat-inactivated at 56 °C for 30 minutes and diluted in DMEM (1:16, 1:32, 1:80, 1:160, 1:320, 1:640, 1:1280). Each dilution was incubated with 20-30 PFU of SARS-CoV-2 at 37 °C for 1 hour. The virus-antibody complexes were added to Vero E6 cells in a 6-well plate at 37 °C for 1 hour. The PFU assay was performed as described above. The PRNT_90_ was calculated as the reciprocal of serum dilution, which inhibited the number of plaques by 90% compared to virus-DMEM control without antibodies ^60^.

### 4.8. RNA isolation from hamster lungs

Total RNA was extracted from the lungs of uninfected and SARS-CoV-2 infected hamsters with or without immune suppression using Trizol reagent and purified by RNeasy mini columns (Qiagen, CA, USA) as described previously ^61^. cDNA synthesis was performed using a High-Capacity cDNA Reverse Transcription Kit with 500ng of total RNA, as per the standard protocol (Applied Biosystems, CA, USA).

### 4.9. RNAseq analysis of lung transcriptome

The quality of RNA was checked for integrity on an Agilent 2200 Tape Station (Agilent Technologies, CA, USA), and samples with RNA integrity number (RIN) >7.0 were used for subsequent processing. Total RNA was subjected to two rounds of poly(A) selection using oligo-d(T)25 magnetic beads (New England Biolabs, CA, USA). Illumina compatible RNAseq library was prepared using NEB next ultra RNAseq library preparation kit. The cDNA libraries were purified using AmpureXP beads and quantified on an Agilent TapeStation and Qubit 4 Fluorometer (Thermo Fisher Scientific, CA, USA). An equimolar amount of barcoded libraries were pooled and sequenced on the Illumina NovaSeq platform (Illumina, San Diego, CA) using the 1×100 cycles configuration. CLC Genomics Workbench 20.0.4 version (Qiagen, Valencia, CA, USA). De-multiplexed fastq files from RNA-Seq libraries were imported into the CLC software. Bases with low quality were trimmed, and reads were mapped to reference genome *Mesocricetusauratus* (assembly BCM_Maur_2.0). The aligned reads were obtained using the RNA-Seq Analysis Tool of CLC Genomics Workbench. Statistical analysis of differentially expressed genes was carried out based on a negative binomial model using the CLC Genomic Workbench tool. Replicates were averaged and statistically differentially expressed genes (SDEG) were identified with a false discovery rate (FDR) p-value <0.05 and fold change of an absolute value >1.5. We compared the following groups: 1) immunocompetent uninfected versus SARS-CoV-2 infected animals at 4 dpi and 16 dpi; (2). Immunosuppressed, SARS-CoV-2 infected animals at 4 dpi and 16 dpi versus immunocompetent animals at respective time points. To account for the gene expression changes caused by immunosuppression treatment, the data from the immunosuppressed/infected animals were normalized to the corresponding uninfected (immunosuppressed) controls before comparison analysis with the corresponding immunocompetent counterparts.

### 4.10. Gene network and pathway analysis

The SDEG was subjected to further visualization analysis, including heat map using Partek Genomics Suite version 7.0 (Partek Inc., St. Louis, MO) software as described previously ^61, 62^. The SDEG were also analyzed by using Ingenuity Pathway Analysis (IPA) software (Ingenuity® Systems, Inc. Redwood City, CA) to determine the networks and pathways that are affected in SARS-CoV-2 infected and/or Uninfected hamster lungs with or without immunosuppression at 4 dpi and 16dpi, as described previously ^61, 62^. In this software, the significance of a network or pathway is determined by both the p-value calculated using the right-tailed Fisher’s Exact Test and the Z-score. For these statistical calculations, the total number of genes in the IPA knowledgebase was compared and computed against the experimental data set from each test group.

### 4.11. Determination of total viral load by quantitative PCR

Quantitative PCR was performed using total RNA and SARS-CoV-2 N gene-specific primers (SARS-CoV-2_N-F1: GTGATGCTGCTCTTGCTTTG and SARS-CoV-2_N-R1: GTGACAGTTTGGCCTTGTTG) and Power SYBR Green PCR MasterMix as per the manufacturer’s protocol (Applied Biosystems, CA, USA) ^63^. The N gene-specific primers were used to amplify 97 bp of SARS-CoV-2 N gene by conventional PCR and purified by Qiagen gel extraction kit (Qiagen, CA, USA). The purified N gene PCR products were used in real-time PCR to prepare a standard curve and determine the viral copy numbers in the lung samples.

### 4.12. Statistical analysis

Statistical analysis was performed using GraphPad Prism-8 (GraphPad Software, La Jolla, CA), and the mean ± standard error (SE) values were plotted as graphs. Unpaired Student’s t-test with Welch correction was used to analyze the data between two groups, and one-way ANOVA with Tukey’s correction was used for multiple group comparison. For all the experimental data, p≤ 0.05 was considered statistically significant.

### 4.13. Data availability

All experimental raw data used in this manuscript is available upon request to the communication author. The RNAseq data has been submitted to Gene Expression Omnibus (GEO) of the NCBI (accession number awaited).

## 5. AUTHOR CONTRIBUTIONS

S.S was responsible for conception, supervision, securing funding for the study, data analysis, putting the data together, writing and editing this manuscript. S.R, A.K and R.K. conducted experiments, acquired and analyzed the data. T.C contributed new reagents and helped with data analysis. S.H and P.S. helped in RNA seq data acquisition and analysis. S.R acquired and analyzed the data from virus neutralization and histopathology assays, wrote the draft version, and prepared the figures. All authors have read, edited and agreed for publication.

## ACKNOWLEDGMENTS

The SARS-Related Coronavirus 2, Isolate USA-WA1/2020, NR-52281was deposited by the Centers for Disease Control and Prevention and obtained through BEI Resources, NIAID, NIH. S.R. acknowledges Dr. Pazhanivel Natesan, Ph.D., Professor, Department of Veterinary Pathology, Madras Veterinary College, India, for his assistance with histopathological examination. Dr. Sabbi Lall at Life Science Editors is acknowledged for critical evaluation of the manuscript. This study was supported by a Center for COVID-19 Response and Pandemic Preparedness (CCRP2) grant from Rutgers University to S.S (#302211).

---

**Supplement figure 1.**
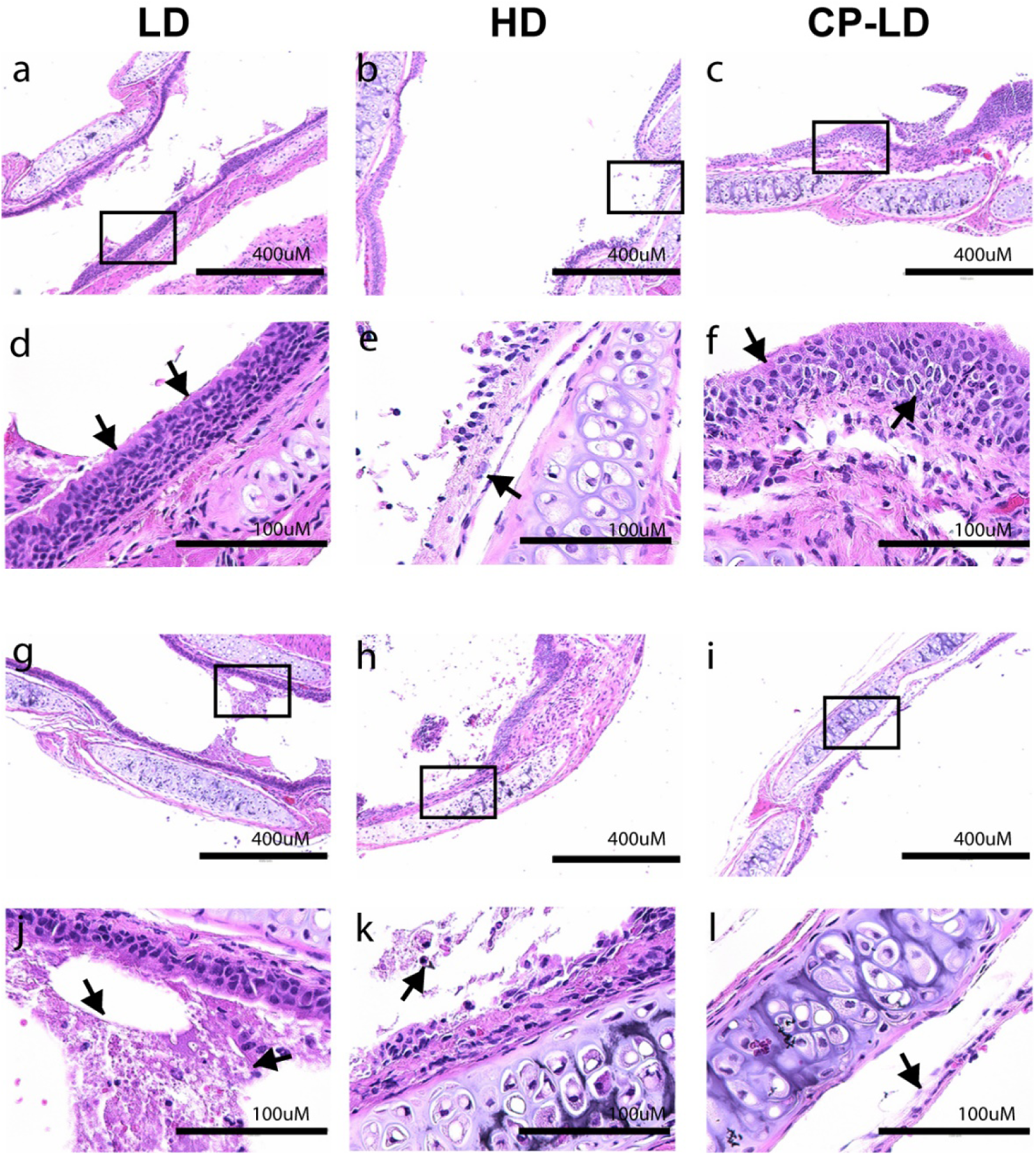
Histopathology of the trachea in SARS-CoV-2 infected hamsters. Histopathological analysis of trachea reveals squamous metaplasia of tracheal epithelial cells (**a**, **b**), mucosal detachment (**b;** *arrow*), severe stratified squamous epithelial metaplasia (**c**) in LD, HD and CP-LD infected hamsters on 4 and 7 dpi. The images in **d**, **e** and **f** area higher magnification of the boxed areas in **a**, **b**, and **c**, respectively. Accumulation of mucinous exudates, macrophages and neutrophils in the tracheal lumen (**g**, **h**). Detachment of epithelial cells (**i, l;** *arrows in* **l**) on 16 dpi in the immunocompromised-LD infected hamsters. The images **a-c**, **g-I** are 100x and **d-f**, **j-l** are 400x magnifications. Scale bar represents 100 µm (**d-f**, **j-l**) or 400 µm (**a-c**, **g-I**). n=3 per time point per group.

**Supplement figure 2.**
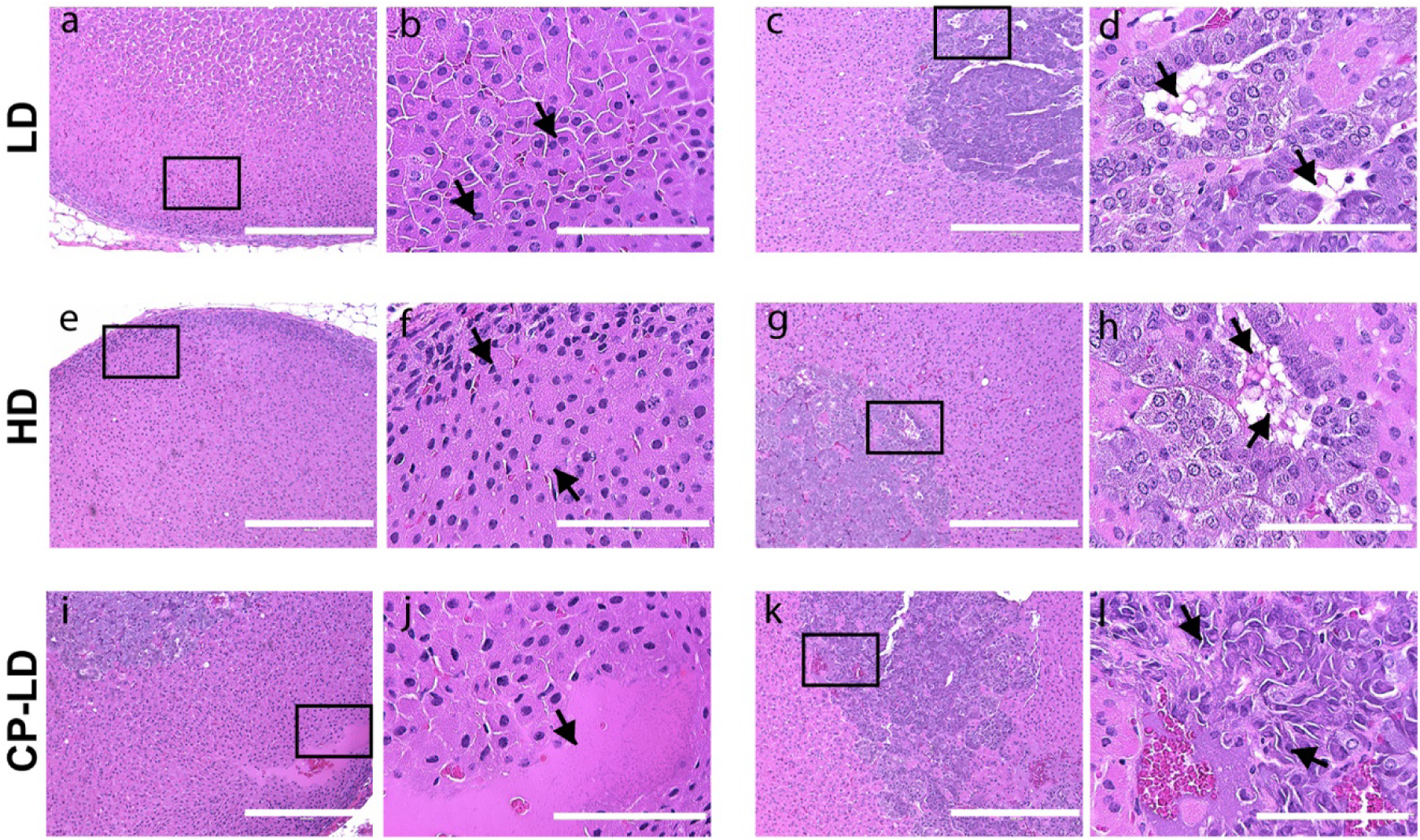
Histopathology of the adrenal gland in SARS-CoV-2 infected hamsters. A section of the adrenal cortex showing degenerated cells with pyknotic nuclei (**a**, **b;** *arrows*) on 4 dpi. and liquefaction necrosis in the medulla (**c**, **d;** *arrows*) on day 7 dpi. Hyperplasia/hypertrophy of adrenal cortex (**e**, **f;** *arrows*) and mild degeneration in the cortical cells (**f**) and liquefaction necrosis in the medullary cords (**g**, **h;** *arrows*) on 7 dpi. In CP-LD infected hamsters, severe degeneration and liquefaction necrosis in the cortex is seen as eosinophilic necrotic material (**i**, **j;** *arrows*) on 4 dpi and chromatolysis and necrosis of chromaffin cells in the medulla (**k**, **l;** *arrows*) on 7 dpi were observed. The images **a**, **e**, **i**, **c**, **g**, **k** are 100x and **b**, **f**, **j**, **d**, **h**, **l** are 400x magnifications (marked in 100x images). Scale bar represents 100 µm (**b**, **f**, **j**, **d**, **h**, **l**) or 400 µm (**a**, **e**, **i**, **c**, **g**, **k**).n=3 per time point per group.

**Supplement figure 3.**
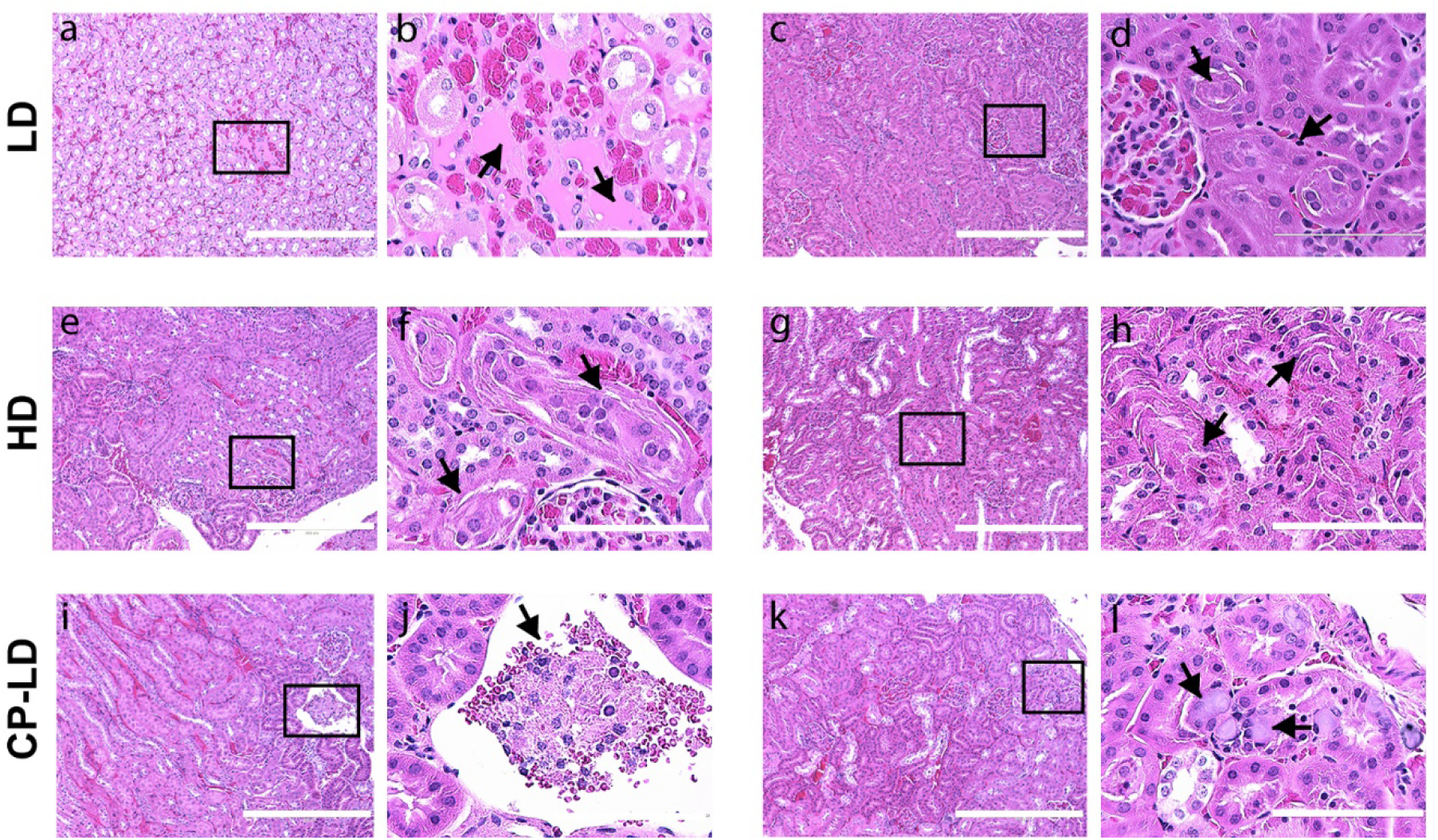
Acute renal tubular necrosis in SARS-CoV-2 infected hamsters. Histopathological analysis of kidney at 4, 7 and 16 dpi revealed severe necrosis of renal tubules with karyolysis and accumulation of eosinophilic material in the tubular lumen (**a**, **b;** *arrows*) and tubular degeneration and mild infiltration of lymphocytes in the interstitium (**c**, **d;** *arrows*) following LD infection (4 dpi). HD infected hamsters’ kidneys showed severe acute tubular necrosis of proximal convoluted tubules with pyknosis and loss of brush border into the lumen (**e**, **f**, **g**, **h;** *arrows*) on 4-16 dpi. Glomerular corpuscular necrosis and detachment from the arterioles (**i**, **j;** *arrow*) and basophilic ground-glass bodies (*arrows)* in the cells of proximal convoluted tubules, pyknosis, and margination of the nucleus (**k**, **l**) in CP-LD infected hamsters on 16 dpi. The images **a**, **e**, **i**, **c**, **g**, **k** are 100x and **b**, **f**, **j**, **d**, **h**, **l** are 400x magnifications (marked in 100x images). Scale bar represents 100 µm (**b**, **f**, **j**, **d**, **h**, **l**) or 400 µm (**a**, **e**, **i**, **c**, **g**, **k**). n=3 per time point per group.

**Supplement figure 4.**
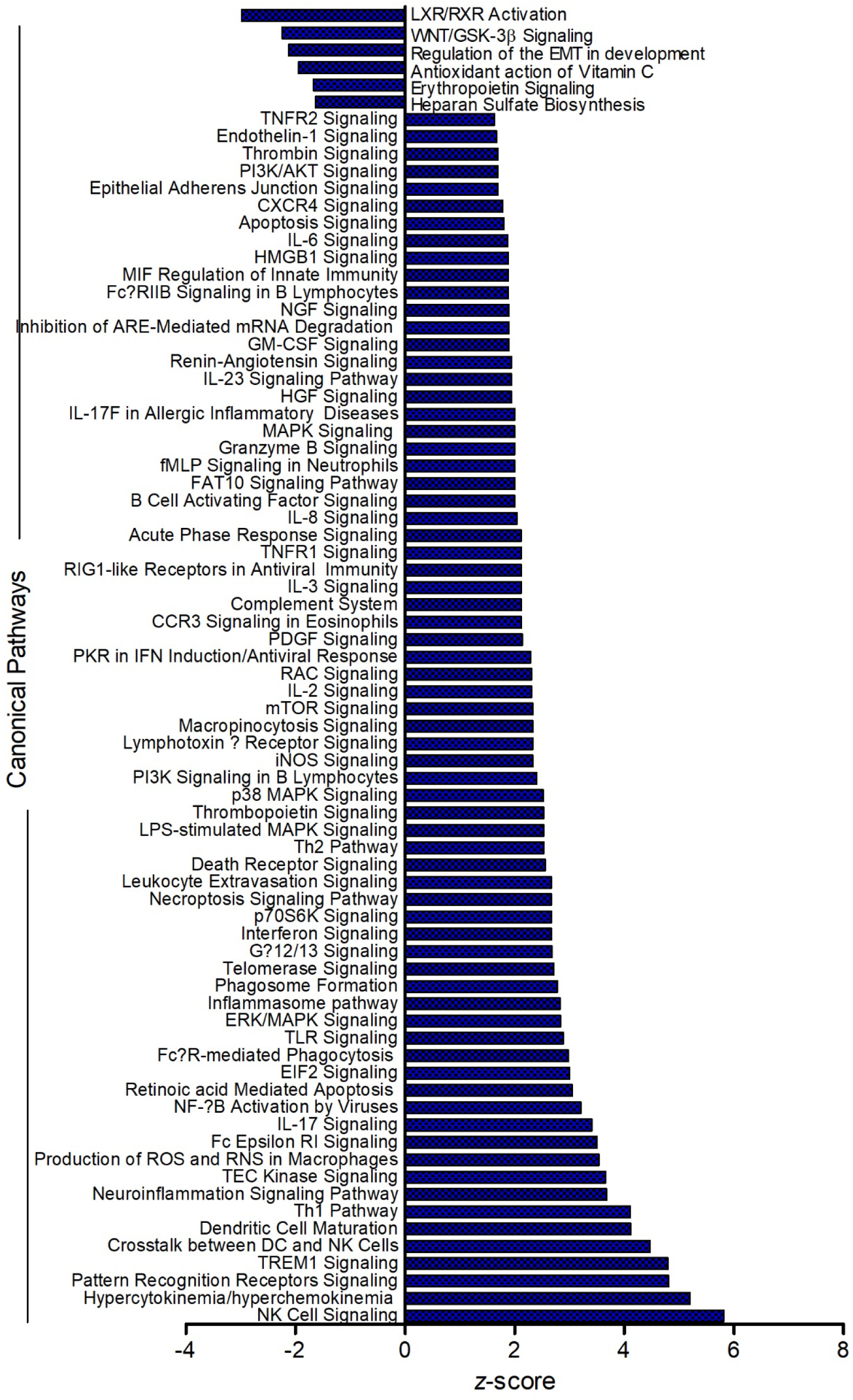
Top canonical pathways operative in immunocompetent hamster lungs infected with SARS-CoV-2 at 4 days post-infection. Pathways with positive z-score values are upregulated/activated, and those with negative z-score values are downregulated/suppressed.

**Supplement figure 5.**
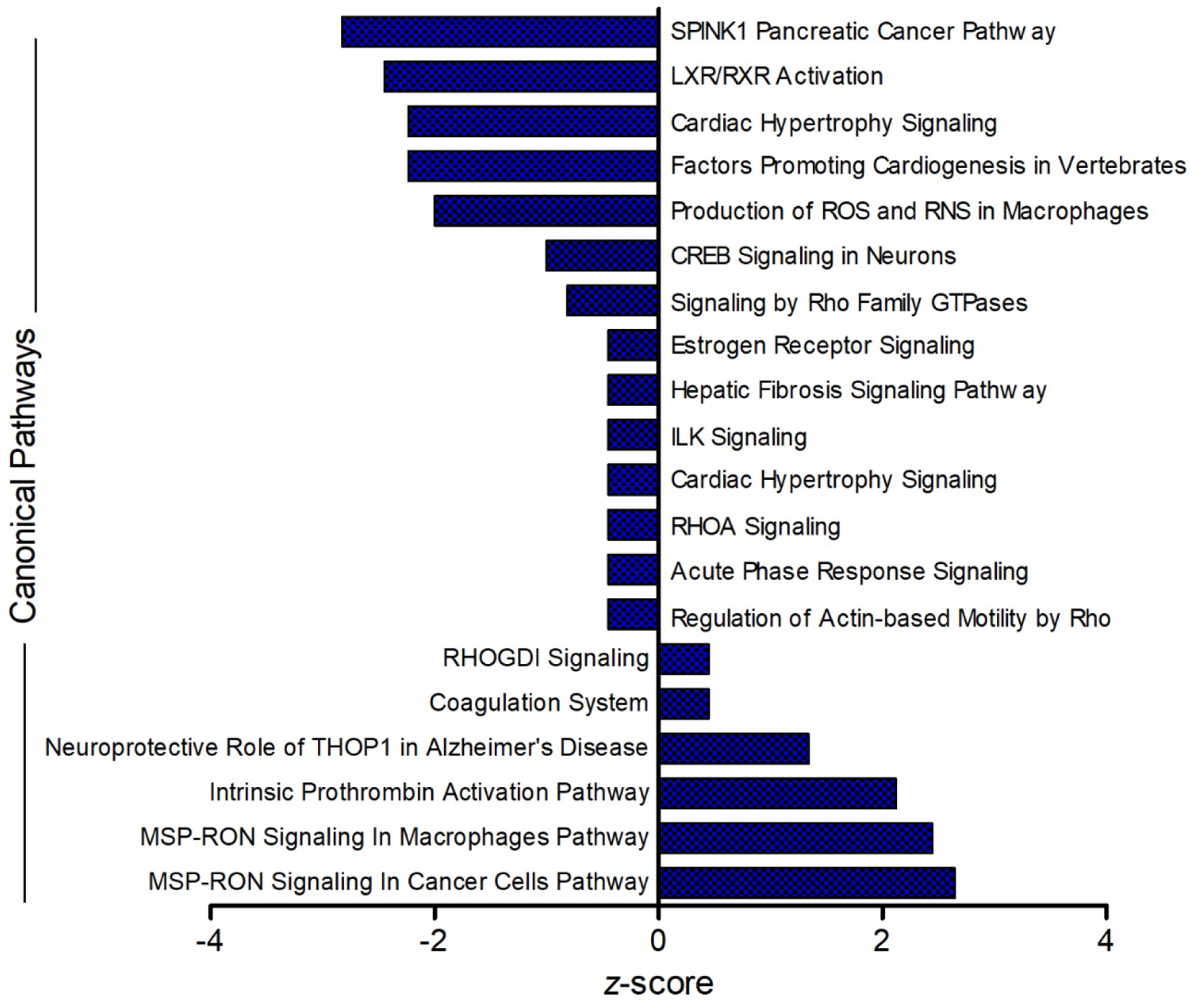
Top canonical pathways operative in immunocompetent hamster lungs infected with SARS-CoV-2 at 16 days post-infection. Pathways with positive z-score values are upregulated/activated, and those with negative z-score values are downregulated/suppressed.

**Supplement figure 6.**
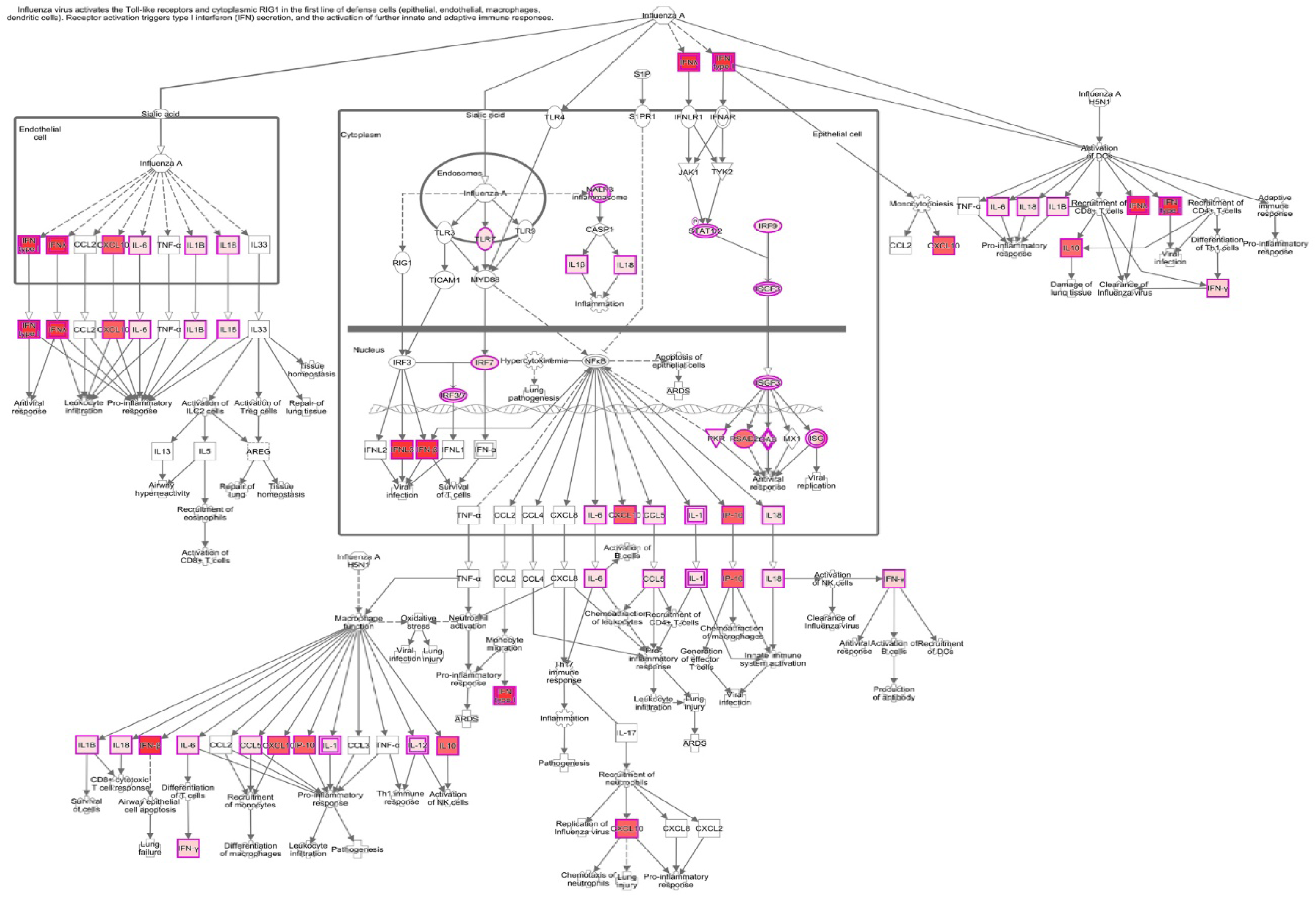
Canonical hypercytokinemia/hyperchemokinemia pathway map. Network genes and their interactions are shown with superimposed data from immunocompetent hamster lungs infected with SARS-CoV-2 at 4 dpi. Red color indicates upregulation, with the intensity of color proportional to the expression level (i.e., darker the color, stronger the expression).

**Supplement figure 7.**
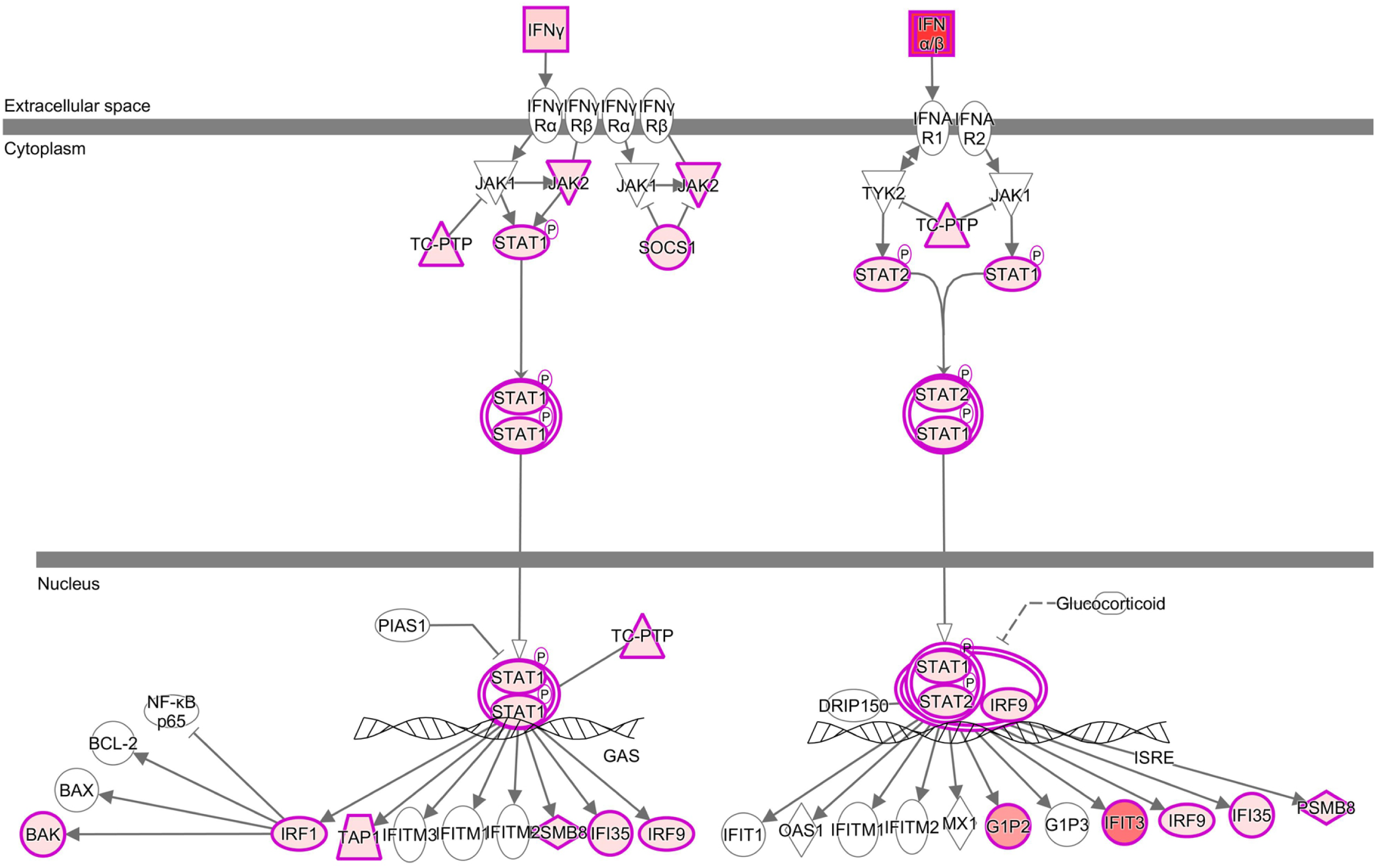
Canonical interferon signaling pathway map. Network genes and their interactions are shown with superimposed data from immunocompetent hamster lungs infected with SARS-CoV-2 at 4 dpi. Red color indicates upregulation, with the intensity of color proportional to the expression level (i.e., darker the color, stronger the expression).

**Supplement figure 8.**
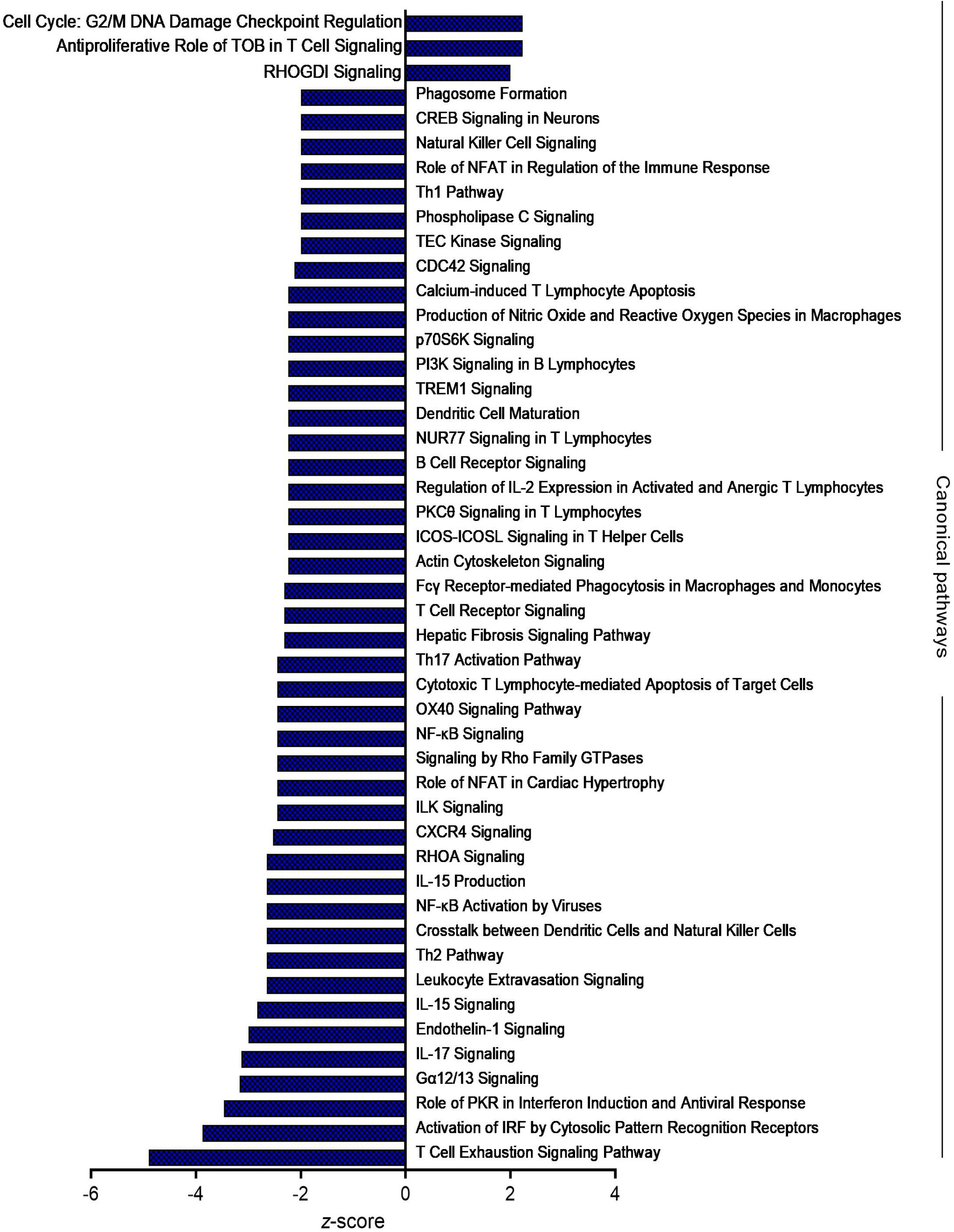
Top canonical pathways operative in immunocompromised hamster lungs infected with SARS-CoV-2 at 4 days post-infection. Pathways with positive z-score values are upregulated/activated, and those with negative z-score values are downregulated/suppressed.

**Supplement figure 9.**
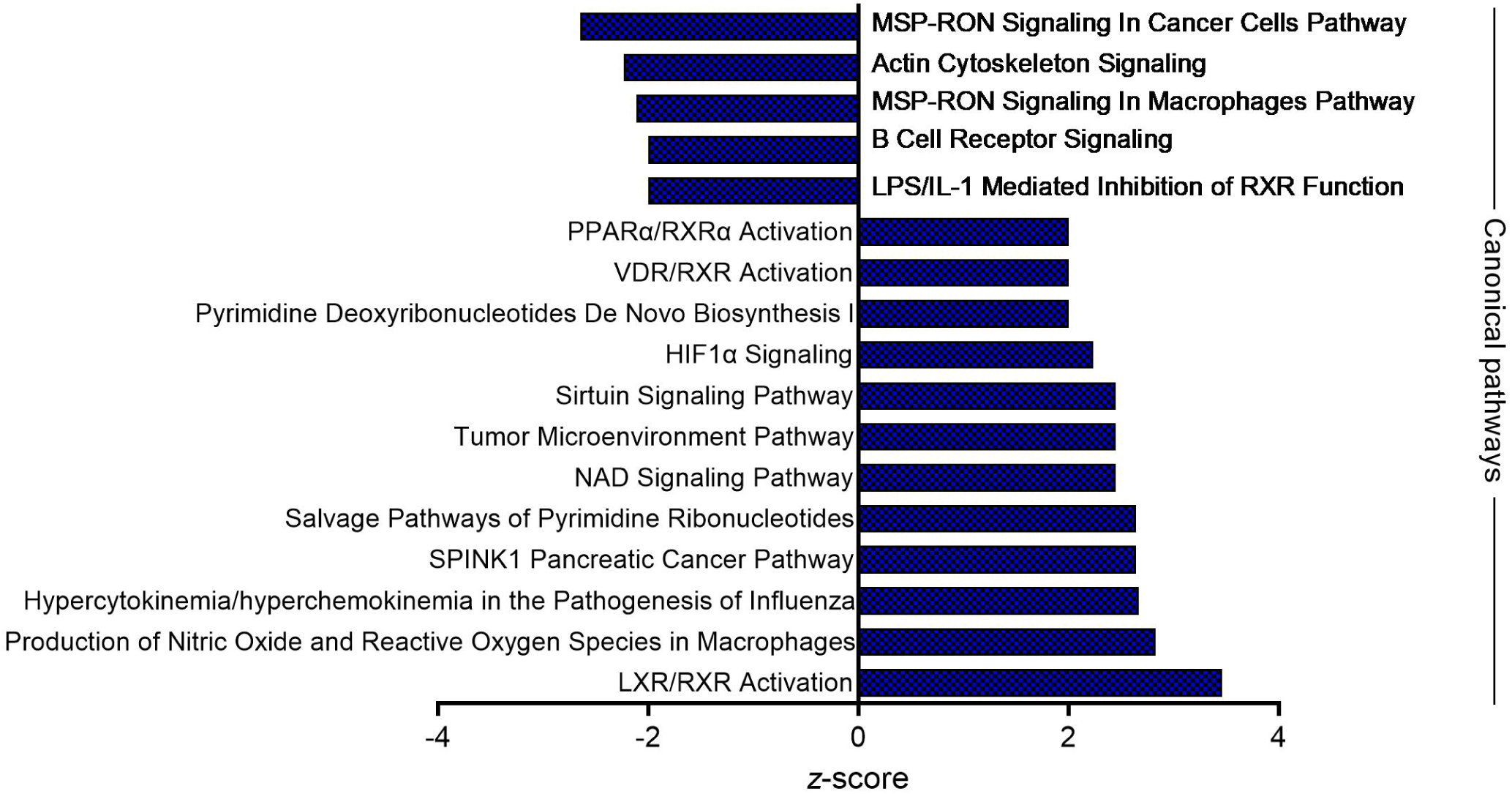
Top canonical pathways operative in immunocompromised hamster lungs infected with SARS-CoV-2 at 16 days post-infection. Pathways with positive z-score values are upregulated/activated and those with negative z-score values are downregulated/suppressed.

**Supplementary Table 1.** Differentially regulated biological functions at 4dpi versus uninfected hamster lungs

| **Categories** | **Diseases or Functions Annotation** | **p-value** | **z-score** | **# Molecules** |
| --- | --- | --- | --- | --- |
| Antimicrobial activities of host cells | Antimicrobial response | 1.74E-34 | 5.136 | 120 |
| Cell Death and Survival | Cytotoxicity | 5.04E-24 | 5.514 | 81 |
|  | Cytolysis | 1.43E-24 | 2.991 | 90 |
| Cell-mediated Immune Response | T cell development | 3.02E-47 | 5.186 | 187 |
|  | Differentiation of T lymphocytes | 6.43E-38 | 5.12 | 139 |
|  | T cell homeostasis | 8.58E-51 | 5.283 | 195 |
|  | T cell migration | 3.84E-34 | 5.073 | 111 |
| Cell-To-Cell Signaling and Interaction | Immune response of myeloid cells | 2.82E-24 | 5.513 | 77 |
|  | Response of antigen presenting cells | 1.11E-26 | 5.295 | 82 |
|  | Binding of myeloid cells | 3.78E-22 | 4.156 | 74 |
| Cellular Growth, Proliferation and Movement | Stimulation of cells | 1.12E-26 | 3.876 | 103 |
|  | Stimulation of lymphatic system cells | 9.23E-25 | 3.409 | 62 |
|  | Stimulation of leukocytes | 8.31E-25 | 4.241 | 70 |
|  | Stimulation of mononuclear leukocytes | 5.29E-25 | 3.65 | 63 |
|  | Stimulation of lymphocytes | 1.51E-25 | 3.487 | 61 |
|  | Stimulation of T lymphocytes | 8.42E-23 | 3.316 | 53 |
|  | Recruitment of blood cells | 1.8E-40 | 5.174 | 130 |
|  | Recruitment of cells | 1.25E-40 | 4.998 | 137 |
|  | Recruitment of myeloid cells | 7.77E-33 | 4.314 | 105 |
|  | Recruitment of leukocytes | 2.36E-40 | 5.192 | 128 |
|  | Recruitment of granulocytes | 1.05E-24 | 4.268 | 79 |
|  | Recruitment of phagocytes | 3.33E-32 | 4.36 | 99 |
|  | Recruitment of neutrophils | 6.83E-24 | 3.993 | 71 |
|  | Interaction of leukocytes | 6.06E-34 | 6.442 | 131 |
|  | Binding of leukocytes | 7.64E-34 | 6.415 | 130 |
|  | Interaction of mononuclear leukocytes | 1.56E-23 | 4.975 | 75 |
|  | Binding of lymphocytes | 9.18E-23 | 4.496 | 63 |
|  | Interaction of phagocytes | 2.21E-24 | 4.483 | 77 |
|  | Activation of myeloid cells | 3.02E-25 | 4.178 | 109 |
|  | Response of lymphocytes | 1.6E-28 | 2.621 | 82 |
|  | Adhesion of immune cells | 3.59E-32 | 6.276 | 121 |
|  | Activation of mononuclear leukocytes | 3.18E-51 | 4.954 | 186 |
|  | Activation of lymphocytes | 2.52E-51 | 4.9 | 181 |
|  | Activation of T lymphocytes | 1.09E-45 | 4.555 | 149 |
|  | Activation of antigen presenting cells | 1.74E-28 | 3.86 | 110 |
| Inflammatory Response | Immune response of leukocytes | 5.24E-41 | 6.073 | 130 |
|  | Immune response of phagocytes | 1.8E-29 | 6.007 | 90 |
|  | Response of phagocytes | 4.06E-31 | 6.004 | 97 |
|  | Immune response of antigen presenting cells | 8.26E-26 | 5.226 | 77 |
|  | Degranulation of myeloid cells | 2.78E-24 | 3.715 | 127 |
|  | Degranulation of leukocytes | 1.07E-28 | 3.556 | 140 |
|  | Degranulation of phagocytes | 2.44E-24 | 3.308 | 126 |
|  | Inflammatory response | 1.14E-63 | 6.031 | 279 |
|  | Cell-mediated response | 2.04E-31 | 3.697 | 85 |
|  | Antibody response | 3.85E-23 | 3.625 | 62 |
|  | Innate immune response | 1.47E-23 | 3.581 | 80 |
|  | Degranulation | 3.13E-35 | 3.572 | 171 |
| Cellular Development | Differentiation of progenitor cells | 4.52E-22 | 3.248 | 129 |
|  | Proliferation of blood cells | 5.84E-67 | 3.852 | 294 |
|  | Development of progenitor cells | 9.63E-23 | 2.914 | 99 |
|  | Differentiation of mononuclear leukocytes | 3.4E-56 | 6.076 | 239 |
|  | Development of mononuclear leukocytes | 2.37E-56 | 5.85 | 239 |
|  | Differentiation of antigen presenting cells | 1.49E-23 | 4.623 | 77 |
|  | Differentiation of phagocytes | 3.91E-24 | 4.441 | 91 |
|  | Development of phagocytes | 4.49E-22 | 3.759 | 83 |
|  | Development of hematopoietic progenitor cells | 3.01E-22 | 2.944 | 94 |
|  | Proliferation of B lymphocytes | 1.13E-23 | 4.06 | 94 |
|  | Proliferation of mononuclear leukocytes | 2.45E-62 | 4.27 | 259 |
|  | Proliferation of immune cells | 5.7E-66 | 4.229 | 274 |
|  | Proliferation of lymphocytes | 3.98E-60 | 4.155 | 253 |
|  | Expansion of T lymphocytes | 6.37E-32 | 3.308 | 74 |
|  | Cell proliferation of T lymphocytes | 2.89E-56 | 3.12 | 217 |
|  | Expansion of mononuclear leukocytes | 2.12E-32 | 2.918 | 83 |
|  | Expansion of lymphocytes | 4.17E-31 | 2.838 | 80 |
|  | Expansion of leukocytes | 7.59E-33 | 2.739 | 87 |
|  | Expansion of lymphoid cells | 1.72E-31 | 2.679 | 81 |
| Cellular Function and Maintenance | Cellular homeostasis | 1.65E-52 | 5.611 | 425 |
|  | Function of leukocytes | 1.35E-76 | 3.069 | 208 |
|  | Function of mononuclear leukocytes | 5.35E-52 | 2.533 | 133 |
|  | Function of lymphocytes | 1.32E-51 | 2.471 | 132 |
|  | Function of T lymphocytes | 1.28E-43 | 2.021 | 109 |
| Free Radical Scavenging | Synthesis of reactive oxygen species | 4.55E-24 | 5.031 | 142 |
|  | Metabolism of reactive oxygen species | 2.73E-24 | 4.901 | 148 |
| Humoral Immune Response,Protein Synthesis | Quantity of IgG | 9.45E-34 | 2.944 | 89 |
| Immunological Disease | Hypersensitive reaction | 3.59E-34 | 3.937 | 143 |
|  | Immunodeficiency | 2.17E-27 | -3.252 | 105 |
| Organismal Survival | Organismal survival | 3.17E-56 | -2.157 | 560 |
|  | Morbidity or mortality | 3.7E-59 | -2.177 | 572 |

**Supplementary Table 2.** Differentially regulated biological functions at 16dpi versus uninfected hamster lungs

| Categories | Diseases or Functions Annotation | p-value | z-score | # Molecules |
| --- | --- | --- | --- | --- |
| Cellular Movement | Cellular infiltration by myeloid cells | 6.41E-06 | 3.241 | 14 |
|  | Cellular infiltration by phagocytes | 2.23E-06 | 2.935 | 14 |
|  | Cellular infiltration by leukocytes | 1.24E-09 | 2.882 | 22 |
|  | Cellular infiltration | 2.22E-09 | 2.871 | 23 |
|  | Cellular infiltration by granulocytes | 7.28E-05 | 2.573 | 10 |
|  | Infiltration by neutrophils | 0.00035 | 2.76 | 8 |
| Hematological System Development and Function | Quantity of macrophages | 0.0004 | 2.532 | 9 |
| Organismal Injury and Abnormalities | Edema | 0.00045 | 2.747 | 12 |
| Organismal Survival and Cell Death | Survival of organism | 1.51E-07 | -3.324 | 27 |
|  | Organismal death | 1.41E-08 | 2.554 | 58 |
|  | Necrosis of muscle | 0.000143 | 2.204 | 12 |
|  | Lung injury | 1.06E-06 | 2.007 | 12 |
|  | Cell death of muscle cells | 0.00045 | 2.033 | 11 |
| Lipid Metabolism, Small Molecule Biochemistry | Fatty acid metabolism | 3.54E-09 | -2.253 | 25 |
|  | Transport of lipid | 0.000118 | -2.381 | 10 |
|  | Export of molecule | 0.000937 | -2.401 | 10 |
|  | Transport of molecule | 0.00097 | -2.613 | 32 |
| Cardiovascular Disease and Abnormalities | Failure of heart | 7.23E-07 | 2.63 | 14 |
|  | Degeneration of heart | 2.16E-09 | 2.599 | 8 |
|  | Congestive heart failure | 1.75E-05 | 2.236 | 8 |

**Supplementary Table 3.** Differentially regulated biological functions in immunosuppressed SARS-CoV-2 infected hamster lungs at 4dpi

| Categories | Diseases or Functions Annotation | p-value | z-score | # Molecules |
| --- | --- | --- | --- | --- |
| Organismal survival | Cell survival | 1.15E-17 | -5.865 | 86 |
|  | Cell viability | 2.72E-16 | -6.371 | 81 |
|  | Cell viability of leukocytes | 1.59E-24 | -4.592 | 38 |
|  | Cell viability of lymphocytes | 8.23E-20 | -4.07 | 29 |
| Cell Morphology | Polarization of helper T lymphocytes | 4.25E-13 | -2.753 | 11 |
|  | Polarization of lymphocytes | 9.9E-15 | -2.495 | 16 |
|  | Polarization of mononuclear leukocytes | 1.33E-15 | -2.645 | 17 |
|  | Polarization of Th1 cells | 6.44E-13 | -2.385 | 8 |
| Cell Signaling and Molecular Transport | Flux of Ca<sup>2+</sup> | 2.37E-16 | -2.614 | 32 |
|  | Mobilization of Ca<sup>2+</sup> | 2.1E-17 | -2.439 | 34 |
| Cell-mediated Immune Response | Differentiation of effector T lymphocytes | 2.47E-11 | -2.056 | 9 |
|  | Differentiation of T lymphocytes | 5.24E-28 | -2.383 | 50 |
|  | Proliferation of thymocytes | 1.91E-14 | -2.802 | 17 |
|  | T cell development | 5.05E-42 | -4.257 | 73 |
|  | T cell homeostasis | 1.93E-44 | -4.457 | 76 |
|  | T cell migration | 4.32E-20 | -2.333 | 36 |
| Cell-To-Cell Signaling and Interaction | Interaction of lymphocytes | 1.83E-16 | -3.367 | 25 |
|  | Interaction of T lymphocytes | 2.17E-14 | -3.125 | 21 |
|  | Activation of lymphocytes | 2.35E-42 | -3.338 | 70 |
|  | Activation of mononuclear leukocytes | 5.82E-42 | -3.428 | 71 |
|  | Activation of natural killer cells | 2.65E-12 | -3.611 | 18 |
|  | Activation of T lymphocytes | 2.23E-32 | -3.199 | 54 |
| Cellular Growth and Development | Maturation of blood cells | 2.92E-14 | -2.455 | 27 |
|  | Maturation of cells | 2.73E-12 | -2.898 | 34 |
|  | Proliferation of blood cells | 6.93E-54 | -2.429 | 104 |
|  | Proliferation of progenitor cells | 5.63E-13 | -3.388 | 31 |
|  | Development of B lymphocytes | 1.54E-13 | -2.097 | 20 |
|  | Proliferation of hematopoietic progenitor cells | 3.24E-15 | -3.216 | 29 |
|  | Dendropoiesis | 1.9E-13 | -2.81 | 20 |
|  | Development of antigen presenting cells | 1.21E-12 | -2.142 | 22 |
|  | Differentiation of antigen presenting cells | 5.42E-16 | -2.836 | 27 |
|  | Hematopoiesis of mononuclear leukocytes | 1.87E-53 | -4.181 | 94 |
|  | Hematopoiesis of phagocytes | 1.19E-14 | -2.075 | 25 |
|  | Leukopoiesis | 3.98E-54 | -4.343 | 101 |
|  | Development of hematopoietic progenitor cells | 5.94E-17 | -2.266 | 33 |
|  | Proliferation of immune cells | 1.83E-56 | -2.465 | 102 |
|  | Proliferation of lymphocytes | 1.9E-57 | -2.301 | 100 |
|  | Proliferation of mononuclear leukocytes | 7.64E-58 | -2.464 | 101 |
|  | Maturation of lymphocytes | 3.47E-16 | -3.003 | 21 |
| Cellular Function and Maintenance | Cellular homeostasis | 9.2E-31 | -4.633 | 111 |
|  | Ion homeostasis of cells | 2.36E-12 | -2.464 | 40 |
|  | Lymphocyte homeostasis | 5.57E-45 | -4.653 | 78 |
|  | Function of helper T lymphocytes | 4.76E-20 | -2 | 20 |
| Cellular Movement and Immune Cell Trafficking | Cell movement of leukocytes | 1.43E-26 | -2.927 | 71 |
|  | Cell movement of lymphocytes | 7.35E-24 | -2.731 | 48 |
|  | Cell movement of mononuclear leukocytes | 4.22E-23 | -2.673 | 51 |
| Hematological System Development and Function | Quantity of B lymphocytes | 5.47E-33 | -2.303 | 54 |
|  | Quantity of CD4+ T-lymphocytes | 4.67E-30 | -2.354 | 41 |
|  | Quantity of lymphocytes | 6.31E-49 | -3.894 | 92 |
|  | Quantity of regulatory T lymphocytes | 6.4E-13 | -2.507 | 19 |
|  | Quantity of T lymphocytes | 2.05E-38 | -3.039 | 71 |
|  | Quantity of leukocytes | 9.58E-51 | -3.909 | 105 |
|  | Quantity of mononuclear leukocytes | 3.42E-48 | -3.771 | 93 |
| Inflammatory Response | Cell-mediated response | 1.1E-17 | -2.29 | 28 |
|  | Inflammatory response | 2.56E-18 | -2.386 | 59 |
| Cell Death and Survival | Apoptosis | 3.45E-22 | 2.226 | 126 |
|  | Apoptosis of blood cells | 7.07E-32 | 2.173 | 60 |
|  | Apoptosis of leukocytes | 2.17E-31 | 2.224 | 57 |
|  | Apoptosis of lymphatic system cells | 2.57E-28 | 3.066 | 49 |
|  | Apoptosis of lymphocytes | 3.78E-28 | 2.963 | 47 |
|  | Apoptosis of mononuclear leukocytes | 2.54E-29 | 2.895 | 49 |
|  | Cell death of lymphatic system cells | 1.62E-33 | 2.698 | 57 |
|  | Cell death of lymphocytes | 1.35E-34 | 2.549 | 56 |
|  | Cell death of mononuclear leukocytes | 7.78E-36 | 2.464 | 58 |
|  | Morbidity or mortality | 5.4E-15 | 5.817 | 107 |
|  | Organismal death | 4.91E-14 | 5.658 | 104 |

**Supplementary Table 4.** Differentially regulated biological functions in immunosuppressed SARS-CoV-2 infected hamster lungs at 16dpi

| Categories | Diseases or Functions Annotation | p-value | z-score | # Molecules |
| --- | --- | --- | --- | --- |
| Antimicrobial and Inflammatory Response | Antimicrobial response | 3.65E-18 | 2.177 | 40 |
| Cellular Movement and Cellular immunity | Cell movement | 8.33E-13 | 2.825 | 118 |
|  | Migration of cells | 1.66E-12 | 2.626 | 109 |
|  | Migration of mononuclear leukocytes | 4.73E-10 | 2.138 | 33 |
|  | T cell migration | 2.47E-09 | 2.223 | 26 |
|  | Cell movement of T lymphocytes | 1.01E-07 | 2.119 | 22 |
|  | Movement of CD4+ T-lymphocytes | 5.62E-07 | 2.295 | 11 |
| Cellular Function and Maintenance | Ion homeostasis of cells | 7.37E-07 | 2.131 | 35 |
| Humoral Immune Response | Quantity of lymphoid tissue | 4.37E-10 | -2.022 | 31 |
|  | Quantity of B lymphocytes | 1.56E-12 | -2.696 | 35 |
|  | Quantity of marginal-zone B lymphocytes | 1.86E-07 | -2.06 | 12 |
|  | Quantity of IgG3 | 1.85E-09 | -2.125 | 13 |
|  | Quantity of IgG2b | 4.97E-08 | -2.234 | 11 |
| Lipid Metabolism and Molecular Transport | Transport of lipid | 4.58E-12 | 3.482 | 27 |
|  | Transport of fatty acid | 6.46E-08 | 2.762 | 11 |
|  | Flux of lipid | 9.23E-08 | 2.519 | 15 |
|  | Transport of steroid | 2.75E-07 | 2.199 | 16 |
|  | Efflux of lipid | 3.47E-07 | 2.359 | 14 |
|  | Efflux of cholesterol | 6.07E-07 | 2.003 | 13 |
|  | Fatty acid metabolism | 9.78E-12 | 2.871 | 45 |
|  | Transport of carboxylic acid | 3.18E-07 | 2.915 | 13 |
|  | Transport of molecule | 1.67E-06 | 3.905 | 72 |
|  | Protein-protein | 6.99E-08 | -2.578 | 18 |
| Organismal Injury and Abnormalities | Fibrosis | 6.92E-14 | -2.792 | 57 |
|  | Morbidity or mortality | 9.64E-12 | -3.901 | 118 |
|  | Organismal survival | 4.96E-11 | -3.853 | 115 |
|  | Thrombus formation | 1.63E-09 | -2.375 | 22 |

